# Mechanistic insights into the allosteric regulation of cell wall hydrolase RipA in *Mycobacterium tuberculosis*

**DOI:** 10.1101/2025.06.28.662095

**Authors:** Giacomo Carloni, Quentin Gaday, Julienne Petit, Mariano Martinez, Daniela Megrian, Adrià Sogues, Mathilde Ben Assaya, Marcell Kakonyi, Ahmed Haouz, Pedro M. Alzari, Anne Marie Wehenkel

## Abstract

D,L-endopeptidase RipA is the major PG hydrolase required for daughter cell separation in *Mycobacterium tuberculosis* (*Mtb*), as RipA defects severely hinder cell division and increase antibiotic vulnerability. Despite extensive studies, the mechanisms governing *Mtb* RipA regulation remain controversial and poorly understood. Here, we report an integrative structural and functional analysis of the SteAB system, a regulatory complex that has been shown to modulate cell separation in the model organism *Corynebacterium glutamicum* (*Cglu*) and is conserved across *Mycobacteriales*. Although *Mtb* SteB was previously proposed to act as a mycobacterial outer membrane copper transporter, the crystal structures of the homodimeric protein, alone and in complex with the RipA coiled-coil (CC) domain, rule out this hypothesis. Instead, the high-affinity SteB-RipA interaction, together with computational and biophysical data, strongly supports the role of SteB as a direct RipA activator that releases enzyme autoinhibition upon complex formation. In addition, crystallographic characterization of the cytoplasmic core of SteA revealed a homodimeric organization harboring a conserved functional pocket similar to the phosphonucleotide-binding site of thiamine pyrophosphokinase. These data, coupled with the *in vivo* phenotypic analysis of a *steAB* knockout mutant of *Cglu*, support a model in which the transmembrane SteAB heterotetramer, driven by cytoplasmic ligand binding, orchestrates the productive periplasmic positioning of RipA, leading to PG hydrolysis activation. These findings shed new light on the regulation of mycobacterial cell wall remodeling, with implications for understanding *Mtb* pathogenesis and identifying novel antimicrobial targets.

## INTRODUCTION

Peptidoglycan (PG), the main component of the bacterial cell wall, consists of glycan strands of two alternating sugar molecules, N-acetylglucosamine (NAG) and N-acetylmuramic acid (NAM), cross-linked by short pentapeptide stems. This cage-like polymer surrounds the plasma membrane, confers cell shape and protection against osmotic disruption, and is continually remodeled during the cell cycle through the coordinated action of different PG hydrolases and synthetases (1). In polar-growing bacteria, such as *Mycobacteriales*, which do not undergo septal constriction during cytokinesis, the PG mesh forms a continuous mechanical linkage between the progeny cells at the nascent division plane. At the end of cell division, this linkage must be released to allow for daughter cell separation (V-snapping) (2). This crucial task is performed by an array of PG hydrolases, whose tight regulation ensures that cell division occurs in a timely and organized manner, preventing uncontrolled cell wall degradation that can lead to morphological defects or bacterial lysis. These defective cell division phenotypes are often accompanied by increased membrane permeability and antibiotic susceptibility, making PG hydrolases attractive targets for the development of novel antimicrobial agents.

In the human pathogen *Mycobacterium tuberculosis* (*Mtb*), D-L endopeptidase RipA (Rv1477) is the major PG hydrolase involved in cell separation (3–5). Although there are other genes encoding PG hydrolases in *Mtb*, RipA is the only endopeptidase whose depletion induces severe morphological defects and *in vivo* reduced infectivity, emphasizing its importance for proper cell separation and integrity. RipA is a member of a conserved clade of the NlpC/P60 enzyme superfamily that cleaves stem peptide bridges within the PG mesh, and has been shown to be important for cell separation in other *Mycobacteriales* species (2, 6–8). Despite its essential role in *Mtb* cell division, limited and controversial information is available regarding the underlying regulatory mechanism(s) of this process. RipA interacts *in vivo* with other cell division proteins, notably the penicillin-binding protein PBP1 and resuscitation-promoting factor RpfB (9, 10), which may be involved in enzyme regulation through protein-protein interactions. Moreover, truncated RipA species were found in cell wall compartments and culture filtrates, suggesting that RipA is proteolytically processed *in vivo* (4), and the protease MarP was reported to hydrolyze RipA during acid stress (11).

More recently, it has been shown that the periplasmic membrane-associated protein SteB binds to and activates full-length RipA in *Corynebacterium glutamicum (Cglu)* by dissociating the intramolecular complex between the catalytic and the EnvC-like coiled-coil (CC) domains (12). SteB is part of a septal transmembrane complex with the cytosolic membrane protein SteA, and inactivation of either *steA* or *steB* (adjacent genes in the same operon) phenocopies RipA inactivation in terms of ethambutol hypersensitivity and cell wall defects (13). These findings demonstrated that the SteAB complex is part of an FtsEX-independent regulatory system for cell wall degradation mediated by RipA (12). However, the underlying molecular mechanisms remain unclear. Although the *steAB* operon is conserved in *Mycobacteriales*, the SteB homolog in *Mtb* (Rv1698) has previously been described as a putative channel-forming protein involved in copper transport (14, 15), and therefore renamed MctB (for Mycobacterial <u>c</u>opper transport protein <u>B</u>). The *Mtb* SteA homolog (Rv1697) is an uncharacterized hypothetical protein annotated as a thiamin pyrophosphokinase that is essential for the *in vitro* growth of *Mtb* H37Rv (16–19). For clarity, we will refer to these proteins respectively as *Mt*SteA (Rv1697) and *Mt*SteB (Rv1698) in the rest of this manuscript.

Here, we report an integrative structural analysis of the SteAB system in *Mtb.* The crystal structures of homodimeric *Mt*SteB, both alone and in complex with the CC domain of RipA, redefine the role of the protein as a regulator of RipA in *Mtb*, ruling out a putative direct role in copper transport. These findings, together with the crystallographic characterization of homodimeric *Mt*SteA, the structural analysis of the *Mtb* SteA/SteB/RipA ternary complex and the phenotypic characterization of a SteAB-defective *Cglu* mutant strain, put forward a model in which, upon cytoplasmic ligand recognition and CC-mediated intracellular signal transduction, formation of the transmembrane SteAB heterotetramer allows for the productive positioning of RipA to activate PG hydrolysis in the periplasm. Elucidating the intricacies of RipA function enhances our understanding of *Mtb* pathogenesis and opens new avenues for the design of innovative strategies to combat tuberculosis.

## RESULTS

### The structures of *M. tuberculosis* SteB, alone and in complex with the RipA coiled-coil domain

*Mt*SteB is a 33 kDa membrane-bound protein (314 amino acids) with a single N-terminal transmembrane (TM) segment. For structural studies, we produced the soluble protein devoid of its TM domain (*Mt*SteB_ΔTM_, residues 38-314) and determined its crystal structure at 2 Å resolution. The protein is a homodimer with a central CC dimerization domain (residues 41-76 from each protomer), surrounded on either side by the respective C-terminal globular cores (residues 77-314) (Fig. 1A). The monomeric core, which displayed a (β/α) topology (Fig. S1A), is similar to that of the homologous SteB from *Cglu* ((12), pdb code 8AU6), with an RMSD of 1.019 Å for 168 Cα equivalent positions (Fig. S1B). A major difference, however, is that *Cg*SteB crystallized as a monomer, whereas *Mt*SteB is a homodimer. The dimeric conformation is mediated by the intermolecular parallel CC formed between the N-terminal α-helices of each protomer (Fig. 1B). The CC was further stabilized by interactions of the helix tip from one protomer with the C-terminus of the second protomer (Fig. S1C). Interestingly, in *Cg*SteB this C-terminal region (residues 296-314) is missing from the construct (12) and could explain why it crystallized as a monomer. Close inspection of the residues involved in CC formation revealed that the heptad repeat pattern is conserved in SteB homologs from other *Mycobacteriales* (Fig. 1C). Despite the lower conservation of the heptad repeat in the *Cglu* protein, the AlphaFold (AF) model of *Cg*SteB predicted the same CC-mediated dimerization mode as seen in *Mt*SteB crystals, thus supporting the SteB homodimer as the functional unit. In the full-length protein, the TM helix immediately precedes the CC helix in full-length SteB (Fig. 1C), and may thus play a role in stabilizing (or modifying) the CC domain structure, suggesting a possible mechanism for conformational signal transduction.

**Figure 1.**
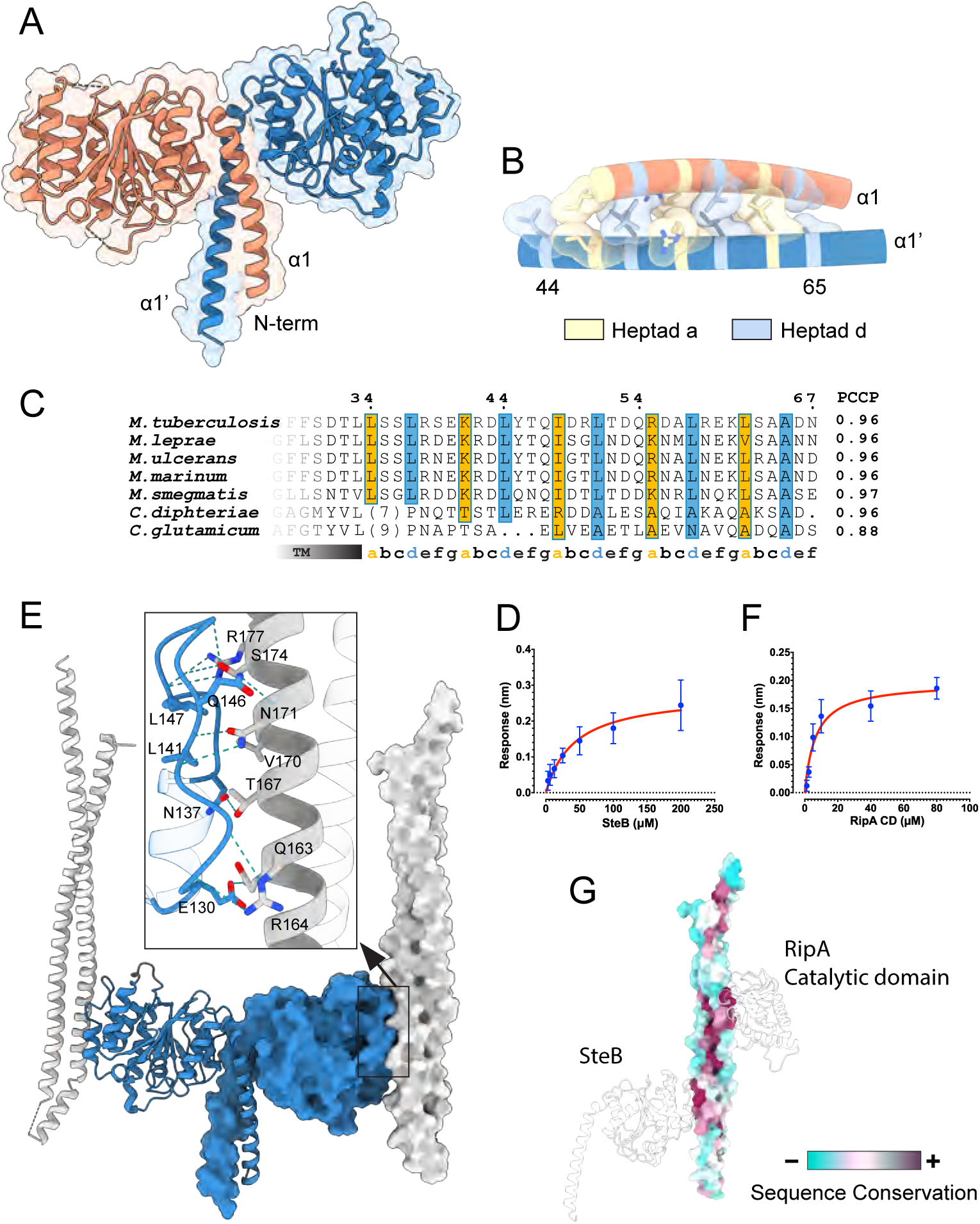
Overall structure of *Mt*SteB. **(A)** Cartoon representation of the SteB homodimer, with each monomer in pink and blue, respectively. **(B)** Detailed view of the N-terminal coiled-coil (CC), showing the residue side-chains at the *a* and *d* positions of the heptad repeats. **(C)** Alignment of the SteB CC region from selected species of *Mycobacteriales*. The TM region and the (*a-g*) heptad repeat positions are shown below the alignment, with the *a* and *d* positions shown in orange and blue respectively. PCCP indicates the probability of parallel CC formation (68). **(D)** BLI binding curves of immobilized *Mt*RipA_CC_ against *Mt*SteB. **(E)** Overall view of the *Mt*SteB_ΔTM_ crystal structure in complex with *Mt*RipA_CC_. The inset shows a ribbon representation of the interface, in which residues involved in hydrogen bonding interactions are labeled. **(F)** BLI binding curves of immobilized *Mt*RipA_CC_ against *Mt*RipA_CAT_. **(G)** The analysis of residue conservation (ConSurf) in all *Mycobacteriales* identifies two non-overlapping regions in the N-terminal CC of RipA (shown here in the same orientation as in panel E) that interact with the C-terminal catalytic domain and the activator protein SteB, respectively.

Given the structural similarity between the SteB homologs from *Mtb* and *Cglu*, it is likely that *Mt*SteB, like *Cg*SteB, plays a direct regulatory role for RipA, rather than the initially proposed role in copper transport (15). Supporting this hypothesis, we observed that *Mt*SteB_ΔTM_ interacted with the CC domain (residues 40-240) of RipA (*Mt*RipA_CC_) with an apparent dissociation constant *Kd* of 42.6 ± 4.4 μM (Figs. 1D and Fig. S2), as determined by bio-layer interferometry (BLI). Furthermore, we crystallized the *Mt*SteB_ΔTM_ *– Mt*RipA_CC_ complex and solved its 3D structure at 2.2 Å resolution (Table S1). The structure revealed a 2:2 heterotetramer (Fig. 1E), where the *Mt*SteB homodimer observed in the *apo* structure was retained. *Mt*RipA_CC_ folds into an antiparallel helical hairpin, with helices α1 (residues 46-120) and α2 (residues 138-238). The two *Mt*RipA_CC_ molecules in the complex aligned parallel to each other and were roughly perpendicular to the membrane plane. Protein-protein association is mediated by hydrogen bonding and hydrophobic interactions between helix α2 of *Mt*RipA_CC_ and the loop connecting helices α3-α4 (residues 139-153) of *Mt*SteB (Fig. 1E inset). Interestingly, the α3-α4 loop, which is well defined in the structure of the complex, was disordered in the *apo Mt*SteB structure and was not visible in the electron density map (Fig. S3), indicating an induced-fit mechanism for RipA recruitment.

### *Mt*RipA is autoinhibited *via* its N-terminal CC domain

The SteB-RipA complex in *Mtb* has an interface similar to that previously predicted for the homologous complex in *Cglu* and was validated by site-directed mutagenesis (12). However, full-length *Cg*RipA crystallized in an autoinhibited form, with its N-terminal CC domain bound to and blocking the catalytic site (12). In contrast, two independent crystal structures of the *Mt*RipA catalytic domain (*Mt*RipA_CAT_), a form that lacks the N-terminal CC domain but includes 16 residues immediately preceding the catalytic domain (residues 264-279) (20, 21), showed that the active site was blocked by this region (Fig. S4A). These structures lent credit to the hypothesis that *Mt*RipA is a zymogen and suggested that its N-terminal CC domain may have a different function (22). To investigate this apparent discrepancy, we predicted structural models of truncated and full-length MtRipA forms using AlphaFold (AF). The active site interactions observed in the previous crystal structures (20, 21) were only observed for constructs lacking the N-terminal CC domain. However, in the full-length protein the active site was predicted to bind the CC (Fig. S4B), to form a complex similar to that observed in the crystal structure of *Cg*RipA (12). To experimentally validate this interaction, we produced separate constructs for *Mt*RipA_CC_ (residues 40-240) and *Mt*RipA_CAT_ (residues 261-472) and measured their binding affinity using BLI. The apparent *Kd* value obtained (6.8 ± 1.4 μM, Fig. 1F and Fig. S4C) strongly supports the CC-mediated autoinhibitory model. In summary, the above findings demonstrate that the *Mt*RipA CC domain uses two well-conserved, non-overlapping regions to bind respectively its own catalytic domain and the activator protein *Mt*SteB (Fig. 1G).

### Structural characterization of SteA

*steB* (*rv1698*) and *steA* (*rv1697*) are part of the same conserved operon in *Mycobacteriales*. The corresponding proteins were described as protein partners in *Cglu* (13) and are encoded by a single fused gene in some species. *Mt*SteA is a 43 kDa protein (393 amino acid residues) that consists of an N-terminal cytoplasmic core followed by a TM helix (residues 344-366, *Mt*SteA numbering) and a C-terminal amphipathic helix exposed on the periplasmic side (residues 371-393). For structural studies we produced a truncated soluble construct (residues 12-344, *Mt*SteA_ΔTM_) that forms a dimer in solution (Fig. S5A) and determined its crystal structure at 2 Å resolution (Table S1). The *Mt*SteA_ΔTM_ homodimer exhibits an overall ‘paper boat’ shape (Fig. 2A), in which the monomers fold into a central C-terminal dimerization domain (the sail) connected through a long (31 residues) α-helix (the hull) to the distal N-terminal globular domains. Protein dimerization buries a largely hydrophobic surface of 2070 Å^2^ from the central linker helix and the C-terminal domain of each monomer (accounting for 13% of the total surface area) and is stabilized by several intermolecular hydrogen bonds and two salt bridges involving residues Arg72 and Asp166 from each protomer.

**Figure 2.**
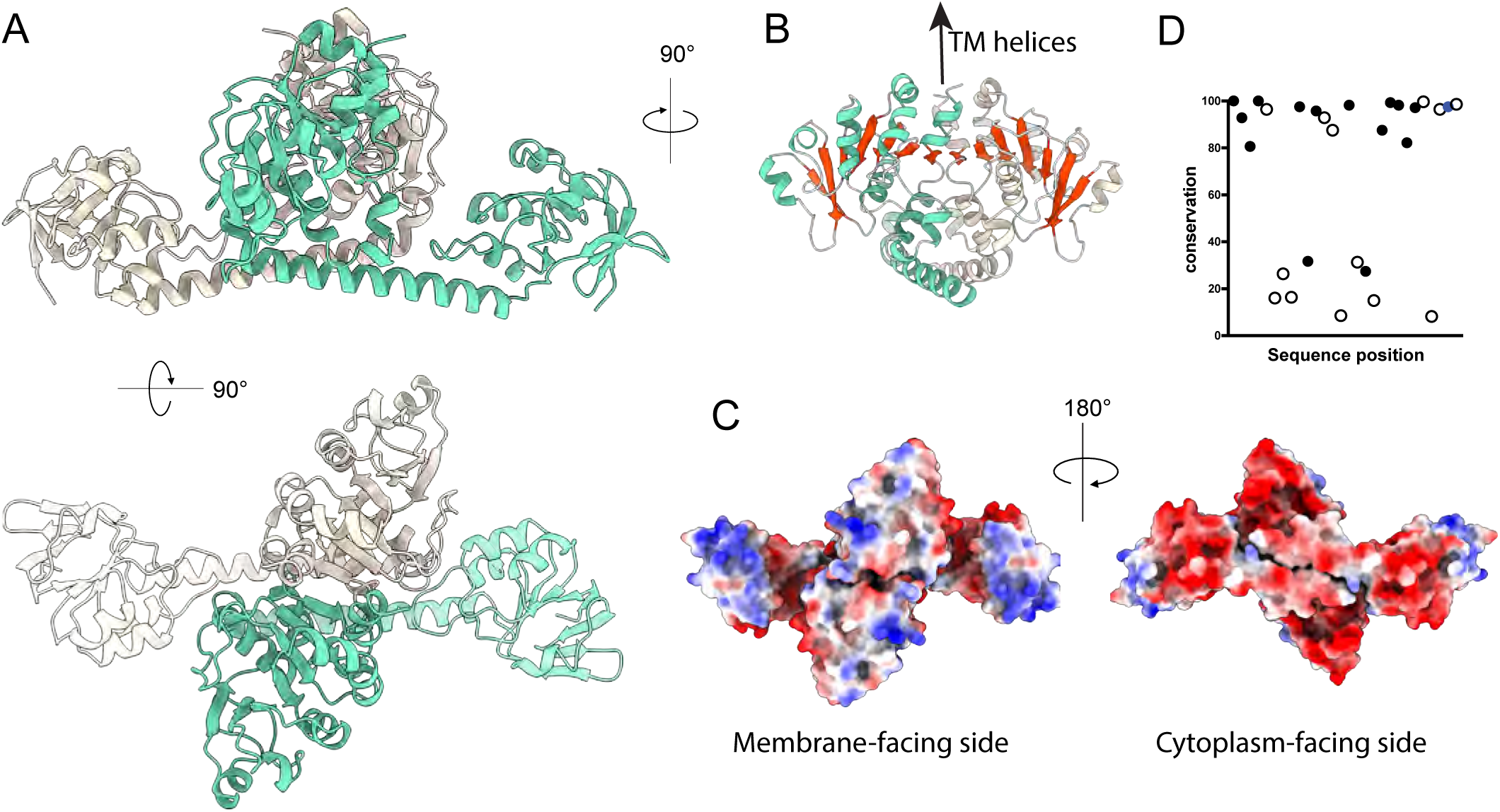
Structure of *Mt*SteA. **(A)** Side and top views of the *Mt*SteA homodimer, with the monomers shown in yellow and green, respectively. **(B)** Protein dimerization results in an intermolecular 14-stranded Δ-sheet (shown in red) formed by the two C-terminal domains. **(C)** Molecular surface of *Mt*SteA colored by electrostatic charges. **(D)** Conservation pattern of all *Mt*SteA basic residues (Arg or Lys) in *Mycobacteriales* SteA homologs. Membrane-facing residue positions, represented by full dots, are generally more conserved.

The N-terminal globular domain (residues 12-127) consists of an external four-stranded antiparallel β-sheet orthogonally packed against a four-stranded parallel β-sheet, covered in turn by three helices (Fig. 2A and Fig. S5B). This three-layer (β/β/⍺) fold resembles that of the swivelling phosphohistidine domain of phosphoenolpyruvate-transferring enzymes (IPR036637) (Fig. S6), although the catalytic histidine is missing in SteA. This structural domain makes only a few intramolecular contacts with the central *Mt*SteA protein core and displays a high intrinsic flexibility. This was confirmed by the crystal structure of the closely similar SteA homolog from *Cglu* (*Cg*SteA), which was determined at 2.05 Å resolution (Table S1) and contained eight independent molecules in the asymmetric unit. In all molecules, the N-terminal domain exhibited high B factors (Fig. S7A), and the overall superposition of the *Cg*SteA and *Mt*SteA monomer structures revealed a wide range of movement of the N-terminal domain (Fig. S7B and Supplementary Movie SM1).

The C-terminal domain of *Mt*SteA (residues 161-342) is responsible for dimerization and forms the central core of the homodimer (Fig. 2A). The domain displayed an (⍺/β)_7_ ovoid fold (Fig. S5B) in which the parallel β-sheet extends, upon dimerization, into a 14-stranded twisted β-sheet flanked by α-helices on both sides (Fig. 2B). At the C-terminus of the dimeric cytoplasmic core, the last α-helices from each protomer immediately preceding the TM helices run parallel to and interact with each other, defining the orientation of the protein with respect to the membrane. In agreement with this hypothesis, the Coulombic electrostatic potential of the protein surface reveals a positively charged membrane-proximal surface (Fig. 2C). This surface includes an array of basic residues (Arg/Lys) that are highly conserved in SteA homologs from *Mycobacteriales* (Fig. 2D) and could interact with negatively charged membrane phospholipids.

Sequence conservation analysis of SteA reveals two clearly distinct patches of conserved residues on the molecular surface of the monomer (Fig. 3A). The larger patch matches the dimerization interface, indicating a conserved homodimerization mode across species. The second patch, on the opposite side of the monomer, corresponds to a protein pocket exposed to the solvent, which may be associated with a ligand-binding site. Structural homology searches using DALI (23) revealed that the C-terminal domain is similar to the N-terminal ATP-binding domain of thiamine pyrophosphokinase (TPPK, (24)) (rmsd of 1.0 Å for 40 equivalent Cα positions) and, to a lesser extent, to sialyl-tranferase CstII from *Campylobacter jejuni* (CstII, (25)) (rmsd of 1.0 Å for 33 equivalent Cα positions). Although SteA homologs are annotated in sequence databases as putative TPPKs because of this partial similarity, the putative thiamine-binding site is completely missing in SteA homologs due to a different quaternary organization of the protein. Instead, the structural superposition of *Mt*SteA with both TPPK and CstII demonstrated that the conserved SteA pocket corresponds to the phosphonucleotide-binding sites for AMP and CMP, respectively (Fig. 3B), strongly arguing for a functional SteA binding site. In our hands, analyzing the binding of various phosphonucleotides to the soluble construct of *Mt*SteA using nano differential scanning fluorimetry (nanoDSF) approaches led to weak, non-specific protein destabilization, which proved inconclusive (Fig. S8). In contrast, a similar investigation on the closely related *Cg*SteA homolog revealed that the addition of GDP or UDP, but not ADP or triphosphate nucleotides, significantly stabilized the protein (Fig. 3C). Although preliminary, these findings suggest that SteA functions as a specific phosphonucleotide-binding protein and imply that cytoplasmic ligand binding and/or hydrolysis might serve as the potential driving force for conformational signal transduction in RipA regulation.

**Figure 3.**
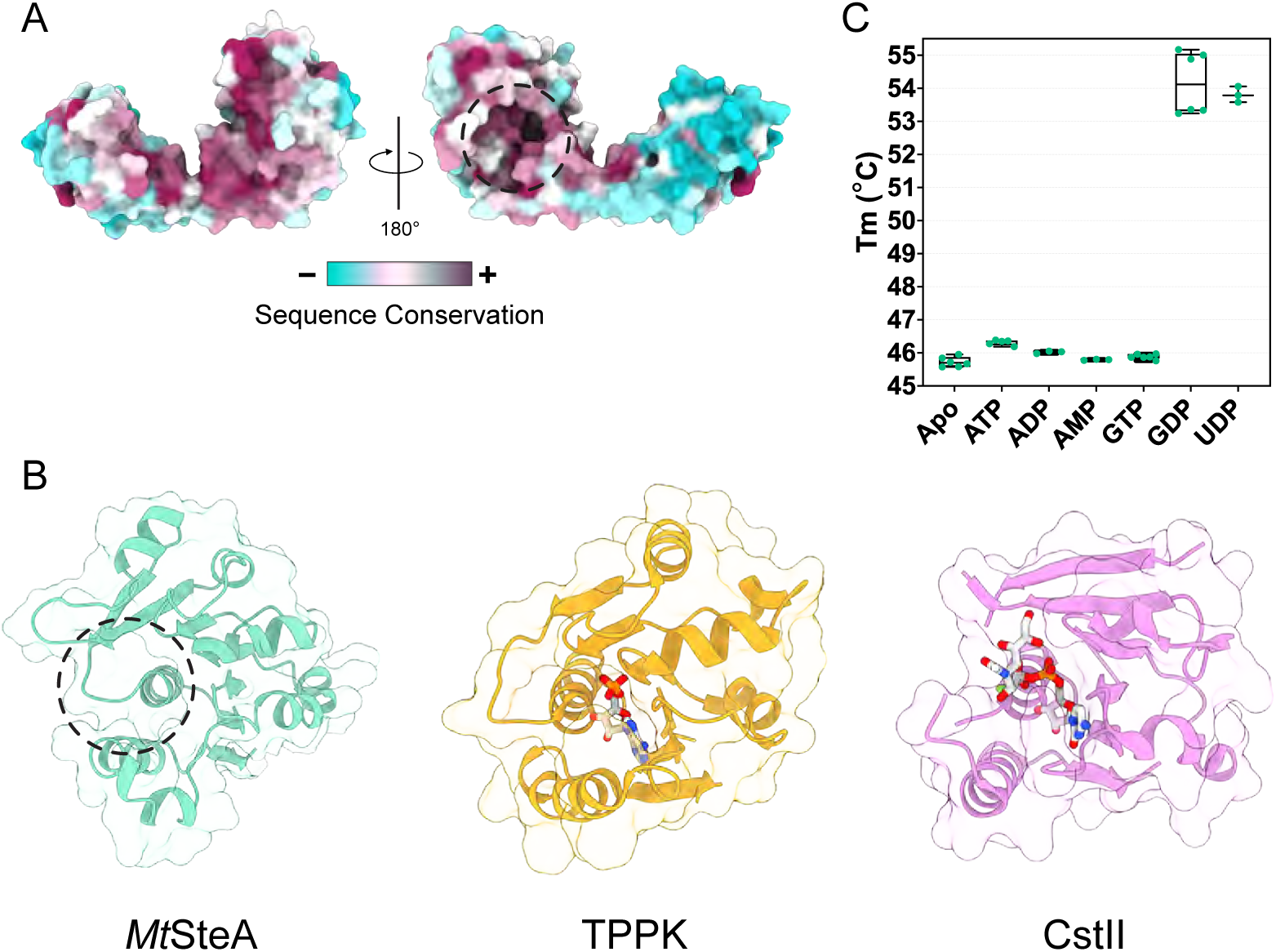
SteA is a putative phosphonucleotide-binding protein. **(A)** Mapping of conserved residues on the molecular surface of the protein, as calculated by Consurf (69). The large, conserved patch in the left panel corresponds to the dimerization interface, and the smaller conserved patch in the right panel corresponds to a putative ligand-binding site**. (B)** The putative *Mt*SteA binding pocket (left) matches the phosphonucleotide-binding sites of TPPK in complex with AMP (PDB 2f17, center) and CstII in complex with CMP (PDB 1ro7, right). The bound ligands are shown in stick representation. **(C)** NanoDSF binding assay of different phosphonucleotides to *Cg*SteA.

### Coordinated and synergistic action of SteA and SteB in cell wall integrity

To understand the role of SteA and SteB *in vivo*, we used homologous recombination (26) to generate a *ΔsteAB* depletion strain in *C. glutamicum* ATCC13032 (*Cglu_ ΔsteAB*). As *steA* and *steB* are the last genes of a predicted 8-gene operon (27) that contains several putative transcription start sites, one of which overlaps with the *steA* coding sequence (Fig. S9A), we hypothesized that the removal of *steA* would also silence *steB* through polar effects. The *Cglu*_ Δ*steAB* strain resulted in elongated, multiseptal cells (Fig. 4A-C) as previously described for the mutants of the individual genes (13). We found that the ectopic expression of *Cg*SteA alone was not sufficient to restore the wild-type *Cglu* phenotype and only expression of both *Cg*SteA and *Cg*SteB fully complemented the mutant strain (Fig. 4A-C). We further confirmed the absence of *Cg*SteB in *Cglu*_Δ*steAB* by anti-SteB Western Blot (Fig. S9B).

**Figure 4.**
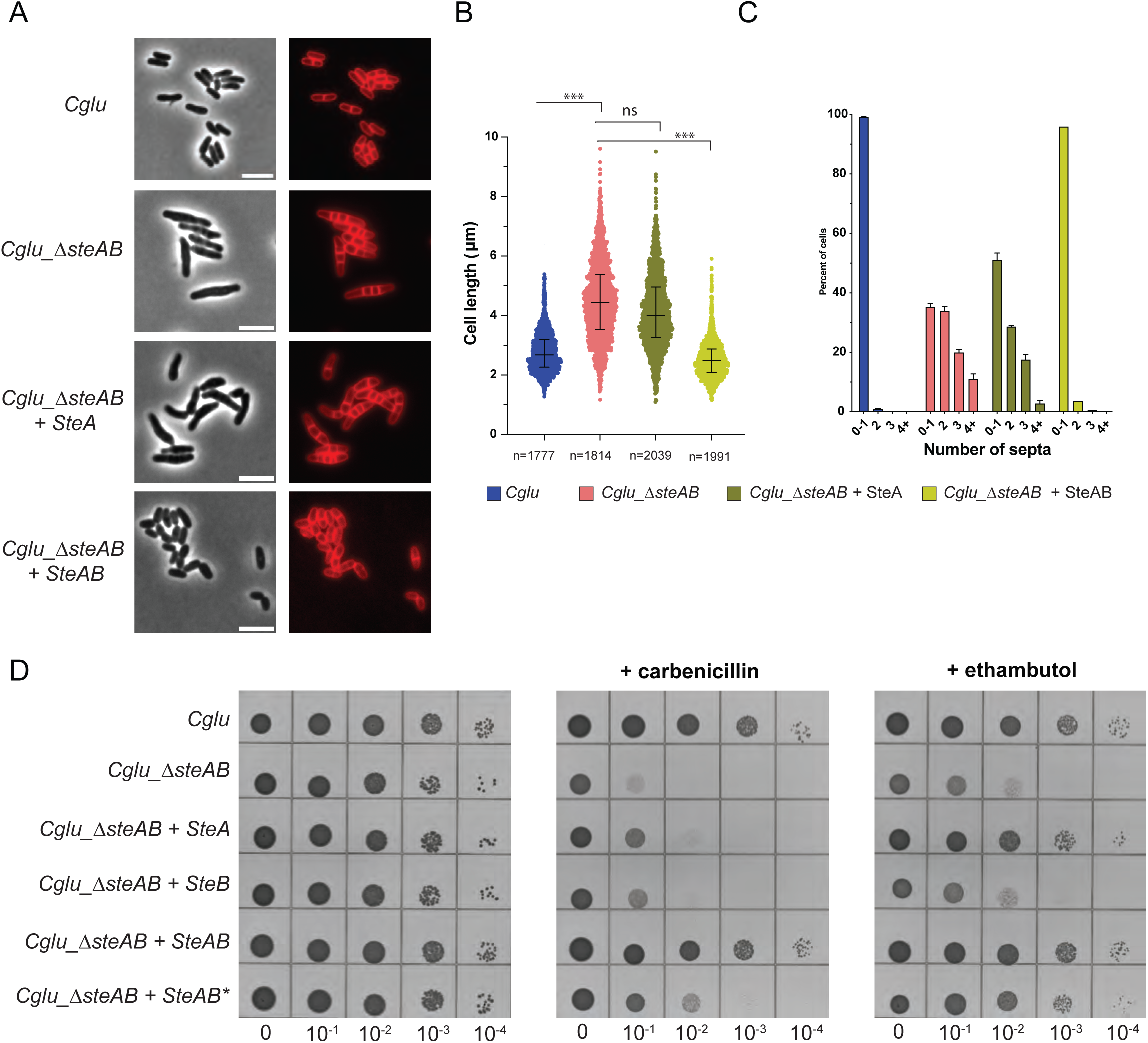
SteA/B depletion and complementation in *C. glutamicum*. **(A)** Representative images in Phase contrast (left) and membrane staining (Nile red, right) for the indicated strains. Scale bars = 5 μm. **(B)** Violin plots showing the distribution of cell length (Cohen’s *d*, from top to bottom: (***, *d* = 1,65, *p* ∼ 0), (ns, *d* = 0,29, *p* = 2,44e-19), (***, *d* = 1,96, *p* ∼ 0)); the whiskers indicate the 25th to the 75th percentile, and the middle line the median. **(C)** Frequency histogram showing the number of septa per cell for the different strains, calculated from 3 independent experiments for each strain. Bars represent the mean ± SD. **(D)** Ethambutol sensitivity assay. BHI overnight cultures of the indicated strains were normalized to an OD600 of 0.5, serially diluted 10-fold, and spotted onto BHI agar medium with or without 1 µg/mL carbenicillin or 0.3 µg/mL ethambutol.

As endopeptidase defects have been associated with an increased sensitivity to cell-wall targeting antibiotics such as β-lactams in *Mycobacteriales* (3, 7), we tested the *Cglu*_Δ*steAB* mutant strain for carbenicillin sensitivity. The mutant displayed antibiotic sensitivity, and wild-type-like carbenicillin resistance could only be restored upon ectopic expression of both *Cg*SteA and *Cg*SteB, but not when expressing the individual proteins (Fig. 4D). To investigate if this phenotype could be due to the SteA/B-mediated control of RipA, we produced the Leu146-Arg point mutant of *Cg*SteB (*Cglu_*SteB_L146R_), that was previously shown to abolish the SteB-RipA interaction *in vitro* (12). We observed that the ectopic expression of *Cg*SteA/*Cg*SteB, but not that of *Cg*SteA/*Cg*SteB_L146R_, restored wild-type-like carbenicillin tolerance in the *Cglu*_Δ*steAB* strain, even if SteB_L146R_ ectopic expression levels were comparable to wild-type *Cg*SteB (Fig. S9B). As *steA* and *steB* were first identified in a transposon mutagenesis screening to identify genes associated with sensitivity to ethambutol (13), we also observed a hypersusceptibility to ethambutol in the *Cglu_ΔsteAB* strain (Fig. 4D). Surprisingly, however, the ectopic expression of *Cg*SteA alone was sufficient to restore wild-type-like ethambutol tolerance (Fig. 4D), suggesting that SteA may have an additional SteB-independent role, possibly mediated by protein-protein interactions with other septal components of the divisome. These findings in *Cglu* could explain why *Mt*SteA, but not *Mt*SteB, is essential for *Mtb* growth (16–19).

### A CC-mediated mechanotransduction mechanism in RipA activation

Previous research in *Cglu* showed that the two transmembrane proteins, SteA and SteB, form a complex that localizes to the cytokinetic ring (13). The AF prediction of this complex in *Mtb* revealed a (*Mt*SteA/*Mt*SteB)_2_ heterotetramer (Fig. S10). On either side of the plasma membrane, SteA and SteB display the same homodimeric arrangement of their soluble cores as seen in their respective crystal structures, and interact with each other through their TM regions, assembled into a four-helix bundle. Integrating this model with the crystal structure of the *MtSteB-MtRipACC* complex described above provides a three-dimensional illustration of the ternary complex involving the SteAB regulatory system and RipA hydrolase (Fig. 5A). In this model, both the SteB CC and the attached RipA CC are oriented at approximately right angles to the membrane plane, positioning the NlpC/P60 catalytic domain deep within the periplasmic space. This configuration enables the catalytic domain to physically interact with the PG substrate and would therefore correspond to the active enzyme state. Furthermore, for the NlpC/P60 domain to perform its role, it must be capable of deeply penetrating the porous PG sacculus, which has an average width of approximately 70 Å in *Mtb* (Fig. 5A). This is achievable because of the relatively long linker (∼100 residues) that connects the NlpC/P60 domain to the *Mt*SteB-bound CC in *Mt*RipA. Although this linker length is well-conserved among mycobacterial RipA homologs, it is consistently longer in corynebacterial homologs (Fig. 5B), matching a thicker PG layer in these species (∼150-180 Å in *Cglu*, Fig. 5A).

**Figure 5.**
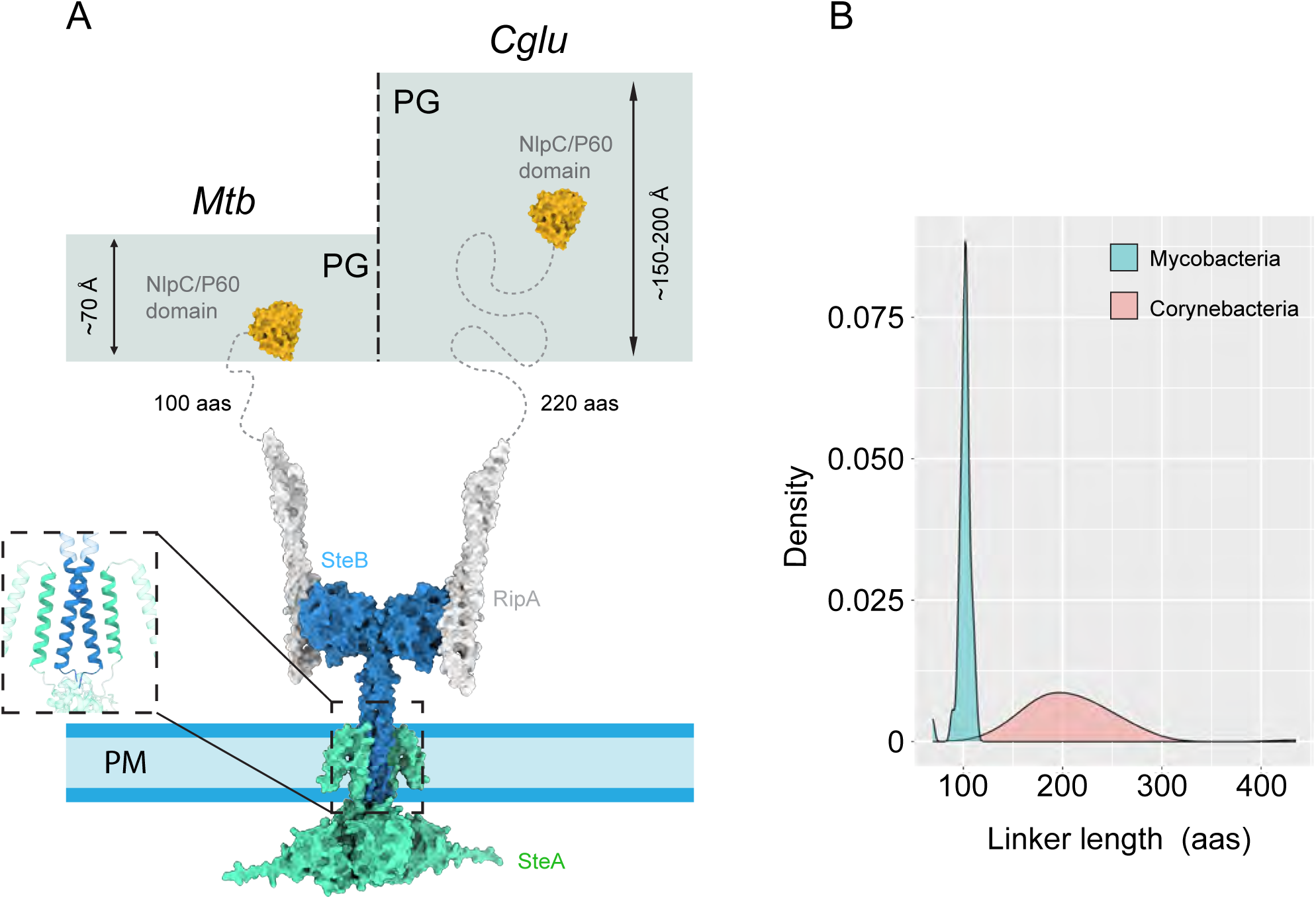
*Mt*SteAB-mediated regulation of *Mt*RipA. **(A)** Proposed model of the ternary complex of SteA, SteB and RipA in an active conformation. The NlpC/P60 catalytic domain (orange) is connected to the N-terminal coiled-coil domain (white) by a flexible linker (dotted line). This connecting linker has different lengths in *Mtb* (shown at left) and *Cglu* (shown at right), which correlates with the approximate thickness of the PG layer in these bacteria (70, 71). The inset shows the predicted TM 4-helix bundle. The inactive state (not shown) might be associated to a SteB conformation with a modified or disrupted CC, as seen for instance for the monomeric *Cg*SteB crystal structure (12), which would preclude the NlpC/P60 catalytic domain from reaching the PG substrate. (**B**) Distribution of connecting linker lengths in RipA homologs from *Mycobacteria* (cyan) and *Corynebacteria* (pink).

## DISCUSSION

The structural characterization of the SteAB complex and its interactions with RipA provides important functional insights into the mechanism by which this system controls daughter cell separation in *Mtb*. The cytoplasmic domain of *Cg*SteA can bind GDP- or UDP-containing molecules, pointing to a possible source of power to trigger SteAB-mediated RipA activation. Although further work is required to identify the specific ligand, different intermediate metabolites and recycling molecules from cell wall synthesis do contain this class of phosphonucleotide moieties (28, 29). This suggests that the system may be able to sense the status of the cell wall to coordinate cytokinesis. Upon cytoplasmic ligand binding (and/or hydrolysis), the initial activation signal is transmitted through the SteAB TM helical bundle to modulate periplasmic RipA recruitment and activation by directly affecting the productive positioning of the catalytic domain for PG hydrolysis. Both four-helix bundles and two-helical CCs are ubiquitous sensory modules involved in bacterial signal transduction (30). Indeed, the proposed SteAB mechanism is reminiscent of those described for bacterial TM histidine kinases, where a conformational signal transmitted along the membrane-connecting two-helical coiled-coil serves as a switching mechanism to control enzyme activity (31–33). As both SteA and SteB have been reported to interact with other divisome proteins (34, 35), we cannot exclude the possibility that additional septal proteins – yet to be identified – could transiently interact with the TM SteAB helical bundle or the cytoplasmic domain of SteA to fine-tune signal transduction.

It was previously proposed that *Mt*SteB (MctB) could be an outer membrane porine involved in copper transport (15). Shortly thereafter, however, the same researchers suggested that the protein might be anchored to the inner membrane and could fulfill a more pleiotropic role (36, 37). Our findings confirm and expand on the second hypothesis. A primary role of *Mt*SteB in the control of daughter cell separation can explain the severe growth defects observed for a ρ*steB Mycobacterium smegmatis* mutant strain or the reduced virulence of a SteB-deficient *Mtb* mutant strain in mice and guinea pigs, originally attributed to copper toxicity (15). Several lines of evidence support the involvement of the SteAB system in cell division: both SteA and SteB were identified as direct or indirect interaction partners of two core divisome proteins, FtsB and FtsQ, in *Mycobacteria* (34, 38); the absence of the cell division membrane protein PerM in a *Mtb* strain resulted in the increased transcription of the two genes, *steA* and *steB* (39); a SteA-deficient *Mycobacterium abscessus* strain displayed a multi-septa phenotype and higher antibiotic susceptibility (40); and transposon insertion mutants of *steA* in *Mycobacterium avium* led to cell wall modifications and reduced multidrug resistance (41).

In addition to the SteAB system, the ABC transporter-like FtsEX is also involved in the regulation of PG hydrolysis in *Mtb* (42). FtsEX controls the action of RipC, another NlpC/P60 hydrolase with a domain organization similar to RipA but whose physiological role remains unclear (43, 44). The recent cryo-EM structure of the FtsEX-RipC complex (45) revealed that RipC binds in an inclined manner and, like RipA, is self-inhibited by its own N-terminal CC domain (Fig. 6). A common feature of these two systems is that enzyme activation relies on the productive periplasmic positioning of the NlpC/P60 domain, as the activation of RipC by FtsEX causes its CC domain to tilt towards the PG layer (45), favoring the physical interaction of the catalytic domain with its substrate. However, these two systems differ markedly in their overall architectures, mechanisms of action and biological roles. FtsEX belongs to subfamily VII of ABC transporters and uses ATP hydrolysis as the power source to control PG hydrolysis. In contrast, SteAB presents a novel architecture, with a cytoplasmic moiety partially resembling TPPKs that binds phosphonucleotide-containing molecules but not ATP, and a periplasmic moiety that constitutively recruits RipA.

**Fig. 6.**
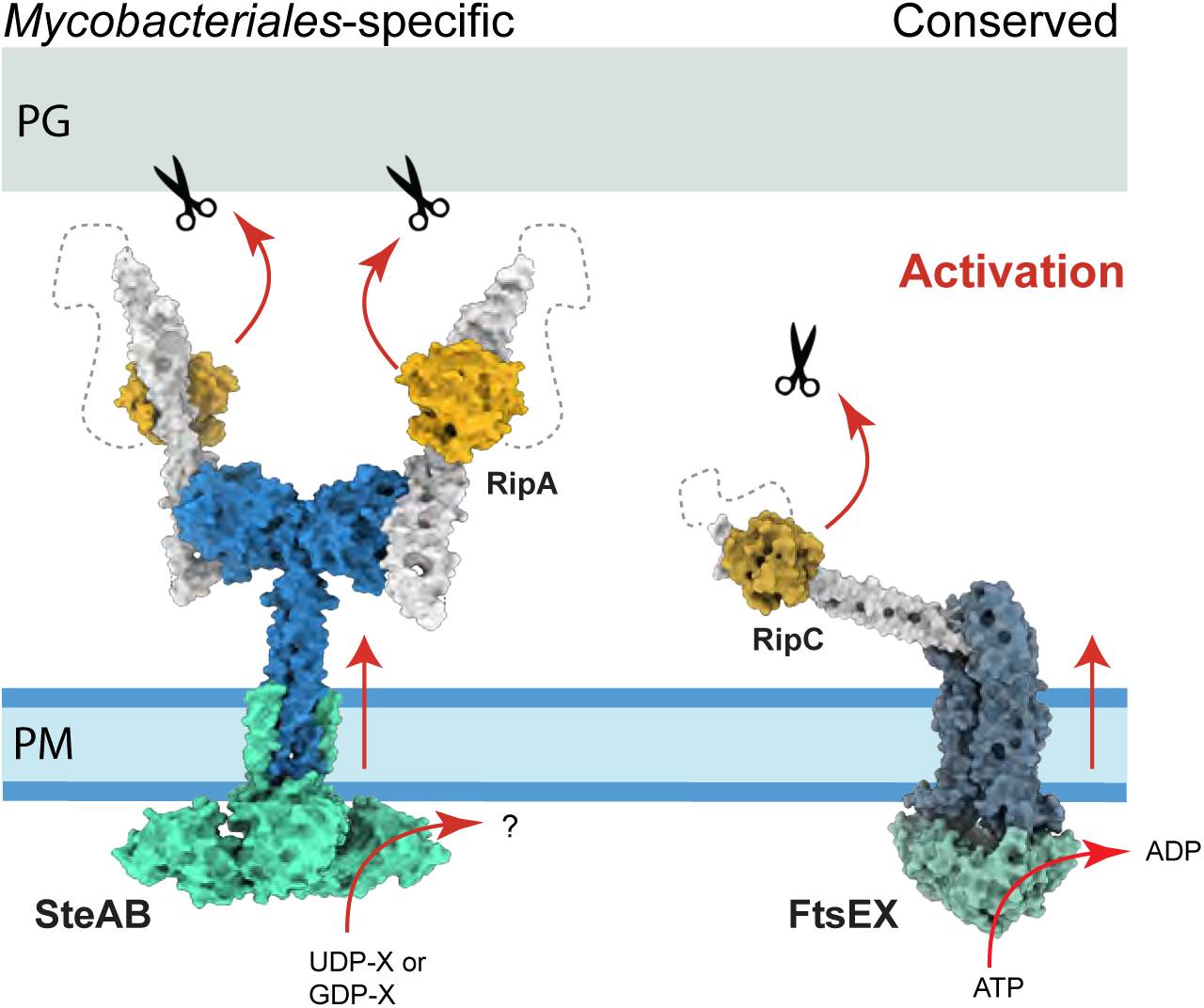
Two regulatory systems of PG hydrolysis in *Mtb*. Overall views of the SteAB-RipA complex (left panel) and the FtsEX-RipC complex (right panel, PDB 8JIA, (45)). The NlpC/P60 catalytic domains (shown in orange) are shown in their autoinhibited state, with their active sites bound to their respective N-terminal coiled-coil domains (in white). In both cases, phosphonucleotide binding to the cytoplasmic domain (SteA or FtsE) would draw the respective NlpC/P60 domains towards the PG layer, where the physical interaction with substrate and/or a conformational signal propagated across the membrane would release the catalytic domain for PG hydrolysis (as illustrated in Fig. 5A for RipA).

FtsEX is highly conserved in bacteria, suggesting an ancient general role in PG remodeling. In contrast, SteAB is largely restricted to *Mycobacteriales*. This group of bacteria has a thick waxy cell envelope formed by the plasma membrane, a two-layered cell wall composed of PG and arabinogalactan, and an unusual outer membrane composed of mycolic acids, the mycomembrane. As the cell envelope is sequentially assembled at the septal junction, the peripheral PG layer remains continuous, acting as a mechanical link that holds the daughter cells together throughout septation (2). The primary role of RipA is to cleave this stress-bearing PG layer, asymmetrically weakening the mechanical strength of the cell envelope (4, 5) to generate a breaking point for V-snapping. This rapid, mechanically driven cell separation promoted by turgor pressure (2, 46) is a common trait in *Mycobacteriales*, but rare in other bacteria (47). Accordingly, RipA deficiency results in linear chains of non-growing cells (4) that can be induced to divide by the local application of external mechanical forces (48). Understanding these specific mechanisms of cell wall remodeling and cytokinesis could offer new attractive therapeutic targets in the context of a critical human pathogen, *M. tuberculosis*.

## MATERIALS AND METHODS

### Bacterial strains and growth conditions

*Escherichia coli* DH5α or CopyCutter EPI400 were used for cloning and grown in Luria-Bertani (LB) broth or agar plates at 37°C supplemented with 50 µg/mL kanamycin or 100 µg/mL carbenicillin when required. For protein production, *E. coli* BL21 (DE3) (for soluble proteins) and C41 (49) (for membrane proteins) cells were grown in 2YT broth supplemented with 50 µg/mL kanamycin or 100 µg/mL carbenicillin at the appropriate temperature. For the *in vivo* experiments, *Cglu ATCC13032* was defined as the wild-type (wt) strain. All *Cglu* strains generated for this study (Table S2) were grown at 30°C with shaking at 120 rpm in brain heart infusion (BHI) medium or CGXII minimal medium supplemented with 4% sucrose (50). When required, the BHI and CGXII media were supplemented with 25 µg/ml kanamycin (BHI_kan_ and CGXII_kan_).

### Knock-out strain generation of *C. glutamicum*

*CgluΔsteAB* was generated using a two-step recombination strategy with the *pk19mobsacB* plasmid to delete the *steA* coding region as described previously (26). Briefly, approximately 600 bp flanking *steA* upstream and downstream of *Cglu* genomic DNA were amplified by PCR using chromosomal DNA of *Cglu* as a template. The PCR fragments were cloned by Gibson assembly into a linearized *pk19mobsacB*. The resulting plasmid was electroporated into *Cglu*. Successful first recombination events were confirmed by PCR and positive colonies were grown overnight in BHI_kan_ medium. The second round of recombination was selected by growth in BHI plates containing 10% (w/v) sucrose. Kanamycin-sensitive colonies were screened by colony PCR to check for *steA* deletion. Positive colonies were verified by sequencing (Eurofins, France).

### Cloning for recombinant protein production

The primers used for PCR amplification of the different fragments or site-directed mutagenesis are listed in Table S3. Cloning was performed by assembling the purified PCR fragments into the specified pET derivative expression vector using the commercially available NEBuilder HiFi DNA Assembly Cloning Kit (New England Biolabs). For soluble protein production, *M. tuberculosis steA, steB* and *ripA* truncations were amplified by PCR using codon-optimized synthetic genes (GenScript) and cloned into a pET vector containing an N-terminal 6xHis-SUMO tag. For full-length protein production*, steA* and *steB* genes were amplified by PCR using genomic DNA of *C. glutamicum* as template and cloned into a pTGR5 shuttle expression vector (51), under the control of a T7 promoter and with or without an N-terminal 6xHis tag. Co-expression vectors were generated by restriction-ligation cloning.

### Soluble protein expression and purification

All constructs were expressed in *E. coli* BL21 (DE3) using an autoinduction method (52). After an initial incubation of 4 h at 37°C, cells were grown for 20 h at 18°C in 2YT medium complemented with autoinduction supplement and 100 μg/mL carbenicillin. Cells were harvested, flash frozen in liquid nitrogen and stored at −20°C. Cell pellets were resuspended in lysis buffer (50 mM Hepes pH 8.0, 500 mM NaCl, 10 mM imidazole, 5% glycerol, 1 mM MgCl_2_, benzonase, lysozyme, 0.25 mM Tris (2-carboxyethyl) phosphine hydrochloride (TCEP), EDTA-free protease inhibitor cocktails (Roche) at 4°C and lysed by sonication. Cell debris were removed by centrifugation (15,000 g) for 15 min at 4°C, and the supernatant loaded onto a Ni-NTA affinity chromatography column (HisTrap FF crude, Cytiva) pre-equilibrated in buffer A (50 mM Hepes pH 8, 500 mM NaCl, 10 mM imidazole, 5% glycerol). His-tagged proteins were eluted with a linear gradient of buffer B (50 mM Hepes pH 8.0, 500 mM NaCl, 0.5 M imidazole). The fractions of interest were pooled and dialyzed in the presence of the SUMO protease at a 1:100 w/w ratio. Dialysis was carried out at 4°C overnight in SEC buffer (25 mM Hepes pH 8.0, 150 mM NaCl). For *Mt*SteA, a higher salt concentration (500 mM NaCl) is necessary to keep the protein soluble. Cleaved His-tags and His-tagged SUMO protease were removed with Ni-NTA agarose resin. The cleaved protein was concentrated and injected onto a Superdex 75 or 200 16/60 size exclusion column (GE Healthcare) pre-equilibrated at 4°C in SEC buffer. The peak corresponding to the protein was concentrated, flash frozen in small aliquots in liquid nitrogen, and stored at −80°C. Protein concentration was determined spectrophotometrically at 280 nm and purity was confirmed by sodium dodecyl sulfate–polyacrylamide gel electrophoresis (SDS-PAGE).

Se-Met-derived *Cg*SteA was expressed in *E. coli* BL21 (DE3) with all media containing 50 µg/mL carbenicillin. Cells were grown for 8 h at 37°C in 2YT medium and inoculated 1:100 in M9 medium (33.7 mM Na_2_HPO_4_-2H_2_O, 22.0 mM KH_2_PO_4_, 8.6 mM NaCl, 9.4 mM NH_4_Cl, 2 mM MgSO_4_, 0.3 mM CaCl_2_ 0.4% (w/v) D-glucose, 3.8 µM thiamin, 4.1 µM biotin). The overnight culture was diluted 1:50 in fresh M9 medium and grown until OD600 = 0.6. The methionine biosynthetic pathway was inhibited by adding lysine, phenylalanine, and threonine at 100 mg/L; isoleucine and valine at 50 mg/L; and selenomethionine at 60 mg/L. Protein expression was induced 30 min after addition of amino acids by adding IPTG to a final concentration of 1 mM, and cells were grown for 20 h at 18°C, harvested, and flash frozen in liquid nitrogen. Protein purification was performed as described above.

### Membrane protein expression and purification

*Cg*SteA and *Cg*SteB-His full-length proteins were recombinantly co-expressed in *E. coli* C41 strain. Cells were grown in 4 L of LB media at 37°C until a OD_600_ of 0.6-0.8 and then expression was induced by addition of 0.5 mM IPTG with an additional incubation of 4 hs at 30°C. Cells were harvested and flash frozen in liquid nitrogen. All following steps were performed at 4°C unless otherwise specified. Cell pellets were resuspended in Lysis buffer (50 mM Hepes pH 7.5, 300 mM NaCl, 5% glycerol, 1 mM MgCl_2_, benzonase, lysozyme, EDTA-free protease inhibitor cocktails (Roche)) and lysed through 3x passages in a CellD press (Constant Systems) at 2.1 kbar. The lysate was cleared by centrifugation (15 min, 15,000 × *g*) and centrifuged in Ti45 tubes for 1 h at 100,000 x *g* in an Optima L-100 XP ultracentrifuge (Beckman Coulter). Pelleted membranes were resuspended with a Dounce homogenizer in Membrane buffer (50 mM Hepes pH 7.5, 500 mM NaCl, 10% glycerol, EDTA-free protease inhibitor cocktail) and solubilized upon addition of an equivalent volume of Membrane buffer containing 2.4% (w/v) DDM for 30 min at room temperature with gentle rotation. 2 mL of Ni-NTA agarose (Qiagen) slurry equilibrated in IMAC A buffer (50 mM Hepes pH = 7.5, 300 mM NaCl, 40 mM imidazole, 5% glycerol, 0.015% DDM) were added to the solubilized membrane fraction. After overnight incubation with gentle rotation, the flow-through was discarded by gravitational flow on a Poly-Prep chromatography column (BioRad), and the resin washed with 25 mL IMAC A buffer. His-tagged proteins were eluted with 5 mL of IMAC B buffer (50 mM Hepes pH = 7.5, 300 mM NaCl, 500 mM imidazole, 2.5% glycerol, 0.015% DDM). The eluate was concentrated with a Vivaspin concentrator (100 MWCO) and loaded onto a Superose 6 Increase 10/300 size exclusion (SEC) column (GE Healthcare) pre-equilibrated at 4°C in SEC buffer (25 mM Hepes pH 7.5, 150 mM NaCl) + 0.015% DDM. The elution fractions were analyzed on SDS-PAGE and the fractions of interest were pooled, concentrated, and subjected to interaction assay with *Cg*RipA immediately (< 24 hours at 4°C).

### Analytical Size Exclusion Chromatography

Protein mixtures were prepared at a final concentration of 50 µM for each protein in SEC buffer, incubated at 4°C for 1 hour, centrifuged (10 mn, 15,000 × *g*), and injected on a Superose 6 Increase 5/150 SEC column (GE Healthcare) pre-equilibrated with SEC buffer at 4°C. Fractions of 100 µL were collected for analysis by SDS-PAGE.

### SEC-SLS

The oligomerization state of SteA was determined by SEC coupled to a triple detection (concentration detector: UV detector, refractometer; SLS 7°, 90°; viscometer) on a Omnisec RESOLVE and REVEAL instrument (Malvern Panalytical). SteA (100 µL sample at 1-5 mg/mL) was centrifuged for 15 min at 27,000 x *g* and injected on a Superdex 75 Increase 10/300 GL column (GE) pre-equilibrated in 25 mM Hepes pH 7.5, 150 mM NaCl at 20°C. External calibration was done by injecting 10 µl bovine serum albumin (BSA) at 18.3 mg/mL. The refractive index, static light scattering, and viscosity measurements were processed to determine the mass average molecular mass and the intrinsic viscosity using the OMNISEC V11.32 software (Malvern Panalytical, UK).

### Protein crystallization and data collection

Screening for initial crystallization conditions was carried out by the sitting drop vapor diffusion method using a MosquitoTM nanoliter-dispensing system (*TTP Labtech, Melbourn, United Kingdom*) and the established protocols the Crystallography Core Facility of the Institut Pasteur (53). Promising hits were then reproduced and optimized manually using the hanging drop vapor diffusion method. All crystallization experiments were carried out at 18°C. *Mt*SteA (5 mg/mL), crystals grew directly in the purification buffer, 0.5 M NaCl, 25 mM Hepes pH 8, without any further manipulation. *Mt*SteB (11.4 mg/mL) crystals were obtained in 0.1 M CdCl_2_, 0.1 M Na Acetate pH 4.6, 30% PEG 400. Crystals of the complex between *Mt*SteB-*Mt*RipA_CC_ were obtained in 0.01 M CoCl_2_, 0.1 M Na Acetate pH 4.6, 1 M Hexane-1,6-diol. An equimolar solution (200 μM, final concentration) of the two proteins was incubated on ice for 30 minutes prior to the crystallization experiment. Crystals of CgSteA (10 mg/mL) were grown in 0.1 M imidazole 8 pH, 0.2 M calcium acetate, 10% w/v PEG 8K, and those of SeMet-derived *Cg*SteA (10 mg/mL) were obtained in 0.1 M Tris pH 8, 0.2 M CaCl_2_, 0.6 M LiCl, 18% PEG 3350. Upon briefly soaking in a cryo-protectant solution containing the mother liquor supplemented by 33% (v/v) glycerol or PEG 400, crystals were flash frozen in liquid nitrogen. Diffraction data were collected at 100K at the Synchrotron facilities Soleil (Saclay, France) or ESRF (Grenoble, France).

### Structure determination and crystallographic refinement

All diffraction data were processed using XDS (54) and Aimless from the CCP4 software suite (55) using the AutoPROC workflow (56). Crystals of the *Mt*SteB-RipA_CC_ complex showed high solvent content (73%) and a strong anisotropy. Anisotropy corrections with STARANISO (57) were applied to the diffraction data from *Cg*SteA and *Mt*SteB-RipA_CC_ crystals.

Structure determination of *Mt*SteB was carried out using cadmium SAD phasing (cadmium was present in the crystallization solution) on a monoclinic crystal form at 2.5 Å resolution. The CRANK2 pipeline (58) within the CCP4 software suite was used to identify 24 cadmium sites and produce a model of the protein with six molecules in the asymmetric unit. This model was used as a search probe to solve the 2 Å resolution orthorhombic *Mt*SteB crystal form, with three independent protein molecules in the asymmetric unit. The structure of the *Mt*SteB-RipA_CC_ complex was solved by molecular replacement using *Mt*SteB as search model. Despite strong anisotropy and high solvent content (>70%), the initial Fourier maps provided enough information to unambiguously build the missing *Mt*RipA_CC_ molecule. The *Mt*SteA structure was solved by molecular replacement using the AlphaFold predicted coordinates of the N and C-terminal domains, separately. 20 residues corresponding to the central alpha-helix linker were removed and manually re-built in Coot to account for the relative flexibility of the two domains. The structure of *Cg*SteA was determined by SAD phasing at 3 Å resolution using the SeMet-labeled protein and refined against a 2 Å dataset from native *Cg*SteA crystals.

All crystal structures underwent extensive iterative cycles of manual model building with COOT (59) and reciprocal space refinement with PHENIX (60) or BUSTER (61). Non crystallographic symmetry (when appropriate) and TLS constraints were applied during refinement. The final refinement statistics are reported in Table S1. All molecular graphics images were generated using ChimeraX (62).

### AlphaFold calculations

Structural predictions were performed on the High-Performance Computing (HPC) Core Facility of the Institut Pasteur computer cluster using a local installation of AlphaFold2 (v2.3.2). Model type ‘monomer’ or ‘multimer’ were used for monomeric and multimeric predictions, respectively. To ensure unbiased predictions for the different RipA forms, structural templates were disabled in these cases. All other parameters were kept as default and amber relaxation was applied to all output models. All models converged to similar conformations and only the best one for each run is shown in Figs. S4 and S10.

### BLI assays

The affinities of purified *Mt*SteB and *Mt*RipA catalytic domain (*Mt*RipA_CAT_, residues 261-472) towards the *Mt*RipA coiled coil domain (*Mt*RipA_CC_, residues 40-240) were assessed in real-time using a bio-layer interferometry Octet-Red384 device (Pall ForteBio) at 25°C. Biotinylated *Mt*RipA_CC_ was diluted at 10 μg/ mL in buffer A (25 mM HEPES pH 7.5, 150 mM NaCl, 5% glycerol) and immobilized on the commercially available Sartorius Streptavidin biosensors for 5 min at 1,000 rpm followed by a washing step in buffer A for 3 min to remove any loosely bound protein. For the biotinylation reaction, 100 μL of recombinant *Mt*RipA_CC_ (25 μM) was incubated with 20 molar excess of EZ-Link NHS-PEG4-Biotin (Thermo Scientific) following supplier instructions.

Empty sensors were used as reference for unspecific binding. *Mt*RipA_CC_-loaded or empty reference sensors were incubated for 5 min at 1,000 rpm in the absence and presence of serially diluted concentrations of *Mt*SteB (200 to 3.125 μM range) or *Mt*RipA_CAT_ (80 to 1.25 μM range) in buffer A supplemented with BSA at 1 mg/mL and 0.05% Tween 20. Specific signals were obtained by double referencing, subtracting both non-specific signals measured on empty sensors and buffer signals on biotinylated *Mt*RipA_CC_-loaded sensors. Two or three independent experiments were performed for *Mt*SteB and *Mt*RipA_CAT_, respectively; and *Kd* values were obtained from steady-state signal versus concentration curves fitted with GraphPad Prism 9 assuming a one-site binding model.

### Nanoscale differential scanning fluorimetry (NanoDSF) assay

Fluorescence measurements were carried out on the Prometheus NT.48 (NanoTemper Technologies) using standard-grade glass capillaries filled with 10–12 μl of *Mt*SteA or *Cg*SteA at 40 μM in 25 mM HEPES, pH 8, 500 mM NaCl and 5% glycerol. Prior to the experiment, protein samples were centrifuged at 15000xg for 15 minutes to remove any large aggregate. Nucleotides stock solutions were prepared in water adjusting the pH to ∼7.5 with diluted NaOH and added at a final concentration of 8 mM. An excitation power of 100% and a temperature ramp from 15 to 95°C with a slope of 1°C/min were used. The Tm was determined in the PR.ThermControl software as the maximum of the first derivative for the ratiometric (F350/F330) melting curves. All melting experiments were performed at least in triplicates.

### Antibiotic susceptibility assay

Overnight cultures of *Cglu* strains were diluted to OD_600_ = 0.5 in fresh BHI_kan_ medium and serial diluted 1:10 in the same medium. 5 µL of each dilution were spotted on BHI_kan_ plates with or without 1 µg/mL carbenicillin or 0.3 µg/mL ethambutol. Plates were imaged after 28 hours of growth at 30°C on a ChemiDoc Imaging System (Bio-Rad).

### Phase contrast and fluorescence microscopy

For imaging, cultures were grown in BHI at 30°C for around 6 hours, pelleted at 5200 x g at RT and inoculated into CGXII, 4% sucrose and kanamycin (25 μg/mL) for overnight growth. The following day cultures were diluted to OD_600_ 1 in CGXII, 4% sucrose (+/− 1% gluconate) and grown for about 7 hours at 30°C to an OD_600_ of about 5 (early exponential phase). For each sample, 100 μL of culture were pelleted and washed twice with fresh medium. For membrane staining, Nile Red (Enzo Life Sciences) was added to the culture (2 μg/ml final concentration) just prior to placing them on 2% agarose pads prepared with the corresponding growth medium. Cells were visualized using a Zeiss Axio Observer Z1 microscope fitted with an Orca Flash 4 V2 sCMOS camera (Hamamatsu) and a Pln-Apo 63X/1.4 oil Ph3 objective. Images were collected using Zen Blue 2.6 (Zeiss) and analyzed using the Fiji software (63), custom trained Omnipose (64) and MicrobeJ (65).

### Statistics and reproducibility

Because of the important number of cells analyzed in each sample, Cohen’s *d* value was used to describe effect sizes between different strains independently of sample size:

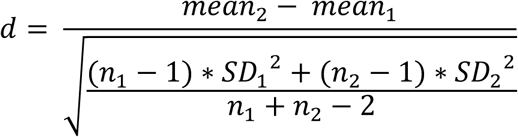

Values were interpreted as previously decribed (66), briefly the intervals of reference are considered: small (ns), *d* < 0.50; medium (*), 0.50 < *d* < 0.80; large (**), 0.80 < *d* <1.20; very large (***), 1.20 < *d* < 2.0; huge (****), *d* > 2.0.

Unless otherwise stated, *p* values were obtained by a Welch two sample t-test calculated on R. All experiments were performed as biological triplicates. Some autofluorescence is observed for wild-*type Cglu* as previously described (67). All micrographs and blots shown are representative of similar experiments carried out at least three times.

### Antibody production and characterization, Western Blots

Polyclonal anti-SteA and anti-SteB antibodies were raised in rabbits (Covalab) against the purified soluble domains of *Cg*SteA and *Cg*SteB respectively. For antibody purification, sera from day 67 post-inoculation were purified using a 1 ml HiTrap NHS-Activated HP column (GE Healthcare) loaded with the corresponding antigen according to manufacturer instructions. Sera were diluted in binding buffer (20 mM Sodium Phosphate pH 7.4, 500 mM NaCl) and loaded onto the column, and washed with 7 ml of binding buffer. Antibodies were eluted with 10 ml elution buffer (100 mM Glycine pH 3, 500 mM NaCl) and neutralized with 1M Tris pH 9. Purified antibodies were concentrated to 8 mg/ml and mixed 1:1 with glycerol 100%, aliquoted and stored at −20°C. The characterization of the antibodies is shown in Fig. S11).

For Western Blots, bacterial pellets of cell extracts were resuspended in lysis buffer (50 mM Bis-Tris pH 7.4; 75 mM 6-Aminocaproic Acid; 1 mM MgSO4; Benzonase and protease Inhibitor) and disrupted at 4°C with 0.1 mm glass beads and using a PRECELLYS 24 homogenizer. Total extracts (from 60 μg to 120 μg) were run on an SDS-PAGE gel, transferred onto a 0,2 μm nitrocellulose membrane and incubated for 1h with blocking buffer (5% skimmed milk, 1X TBS-Tween buffer) at room temperature (RT). Blocked membranes were incubated for 1h at RT with the corresponding primary antibody diluted to the appropriate concentration in blocking buffer. After washing in TBS-Tween buffer, membranes were probed with an anti-rabbit or an anti-mouse horseradish peroxidase-linked secondary antibody (GE healthcare) for 45 minutes. For chemiluminescence detection, membranes were washed with 1X TBS-T and revealed with HRP substrate (Immobilon Forte, Millipore). Images were acquired using the ChemiDoc MP Imaging System (Biorad). Dilutions used: anti-SteA (1:500), anti-SteB (1:1000) and anti-rabbit secondary Abs (1:10000).

## Data availability

Atomic coordinates and structure factors have been deposited in the PDB with accession codes 9HLE (*Mt*SteB), 9HMX (*Mt*SteB-RipA_CC_), 9HMY (*Mt*SteA), 9HMZ (*CgSteA*). All materials of this paper can be provided upon reasonable request.

## Acknowledgements

We gratefully acknowledge the core facilities at the Institut Pasteur C2RT, P. England, B. Raynal, S. Brûlé (PFBMI); P. Weber, C. Pissis, A. Mechaly (PFC), J. Fernandes (UtechS PBI / Imagopole, supported by France BioImaging; ANR-10–INSB–04; Investments for the Future) and the HPC Core Facility of the Institut Pasteur. We also acknowledge the synchrotron sources Soleil (Saint-Aubin, France) and ESRF (Grenoble, France) for granting access to the facilities, and the staff of Proxima 1, Proxima 2A and ID-23-1 beamlines for helpful assistance during X-ray data collection. This work was partially supported by grants from the Agence Nationale de la Recherche (ANR, France), contracts ANR-18-CE11-0017 (P.M.A.), ANR-21-CE11-0003 (A.M.W.), and ANR-21-CE20-0040 (A.M.W), from the Fondation pour la Recherche Médicale (grant number EQU202303016284 to P.M.A.) and by institutional grants from the Institut Pasteur, the CNRS, and Université Paris Cité. For the purpose of open access, the author has applied a CC-BY public copyright licence to any Author Manuscript version arising from this submission. Molecular graphics were done with ChimeraX, developed at UCSF with support from NIH (R01-GM129325) and NIAID.

## Author contributions

A.M.W. and P.M.A. designed the research; G.C., Q.G. and M.K. conducted the protein biochemistry, and purified proteins for structural and biophysical studies; Q.G. and A.S. performed molecular and cell biology and genetic experiments; Q.G. and J.P. performed cellular image analysis; G.C., Q.G. and M.M. and M.B.A carried out the biochemical and biophysical studies of protein-protein interactions; G.C., Q.G. A.H., and P.M.A. carried out the crystallographic and structure prediction studies; G.C., Q.G. and D.M. performed the sequence analyses; A.M.W. and P.M.A. wrote the paper. All authors edited the paper.

## Supplementary information

**Figure S1.**
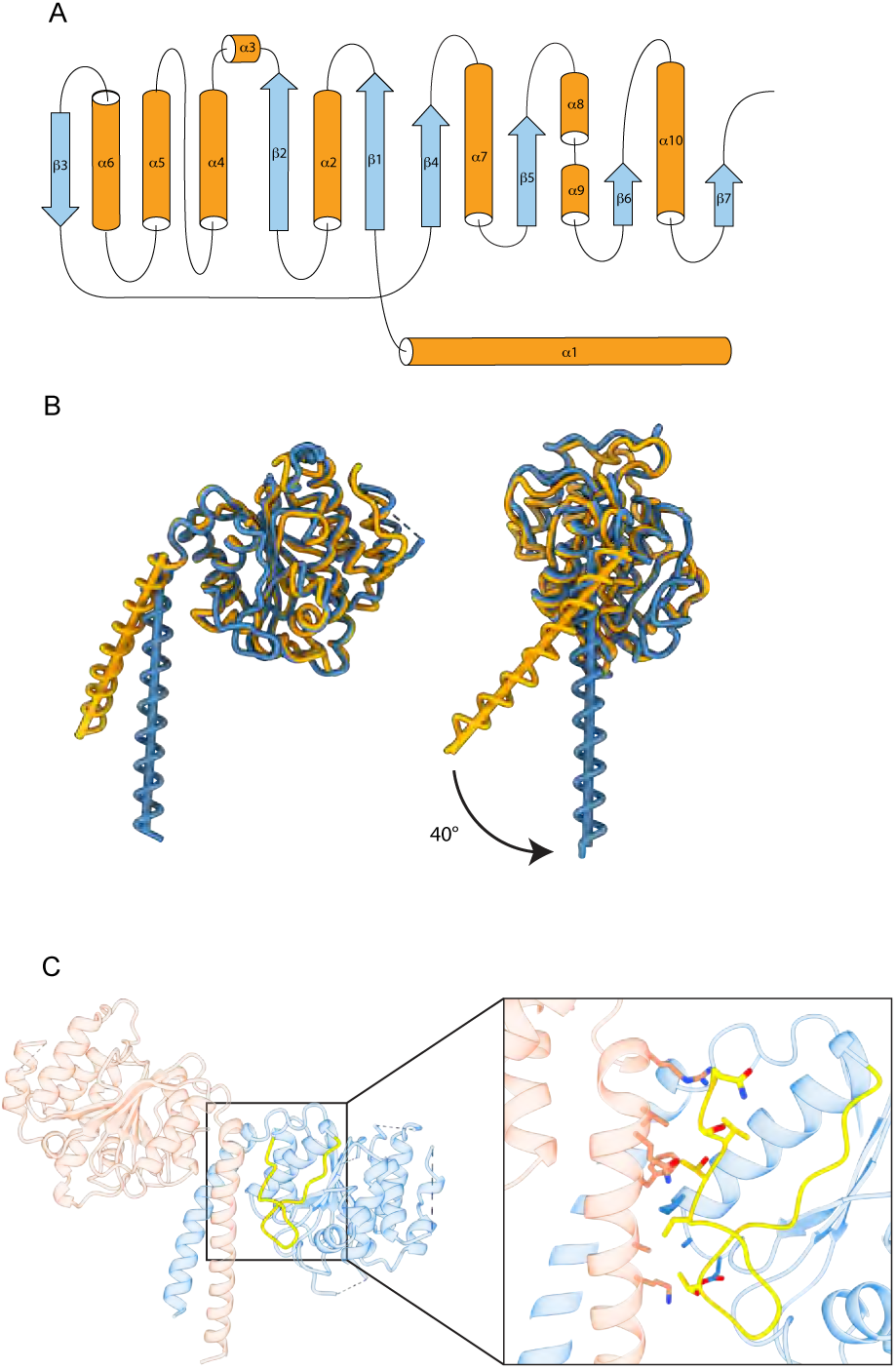
Structure of *Mt*SteB. **(A)** The secondary structure topology of *Mt*SteB partially resembles the (Δ/α)_5_ topology of receiver domains from bacterial response regulators. **(B)** Despite a rather low sequence identity (30%), the structural cores of *Mt*SteB (in blue) and *Cg*SteB (in orange) are closely similar. The major differences are observed in the region connecting the C-terminal globular core with the N-terminal helix, leading to a marked change in the orientation of the major helix axis, as well as at the C-terminus of the protein, which is clearly visible in *Mt*SteB but is absent or structurally disordered in *Cg*SteB. (**C**) Close up view of the dimerization interface between the helix from one protomer with the protein C-terminus from the other. Residues shown in yellow are missing in the construct of monomeric *Cg*SteB (1). The absence of these residues in *Cg*SteB, together with a weaker conservation of the heptad repeat and the lack of the TM region in the construct, might explain why *Cg*SteB crystallized as a monomer.

**Figure S2.**
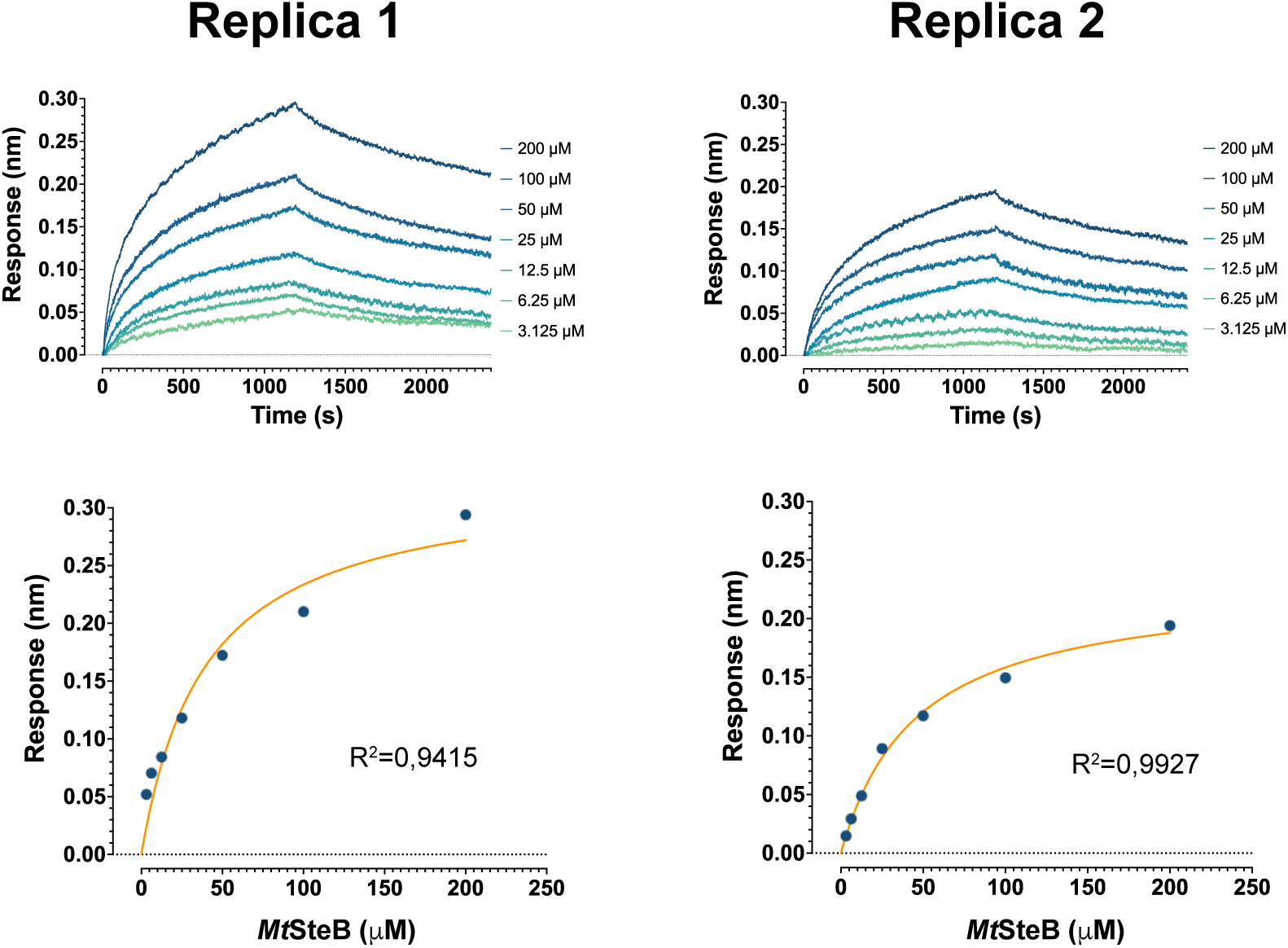
Two independent BLI experiments showing the interaction profiles for the *Mt*RipA-*Mt*SteB interaction used to calculate the dissociation constant shown in Fig. 1D.

**Figure S3.**
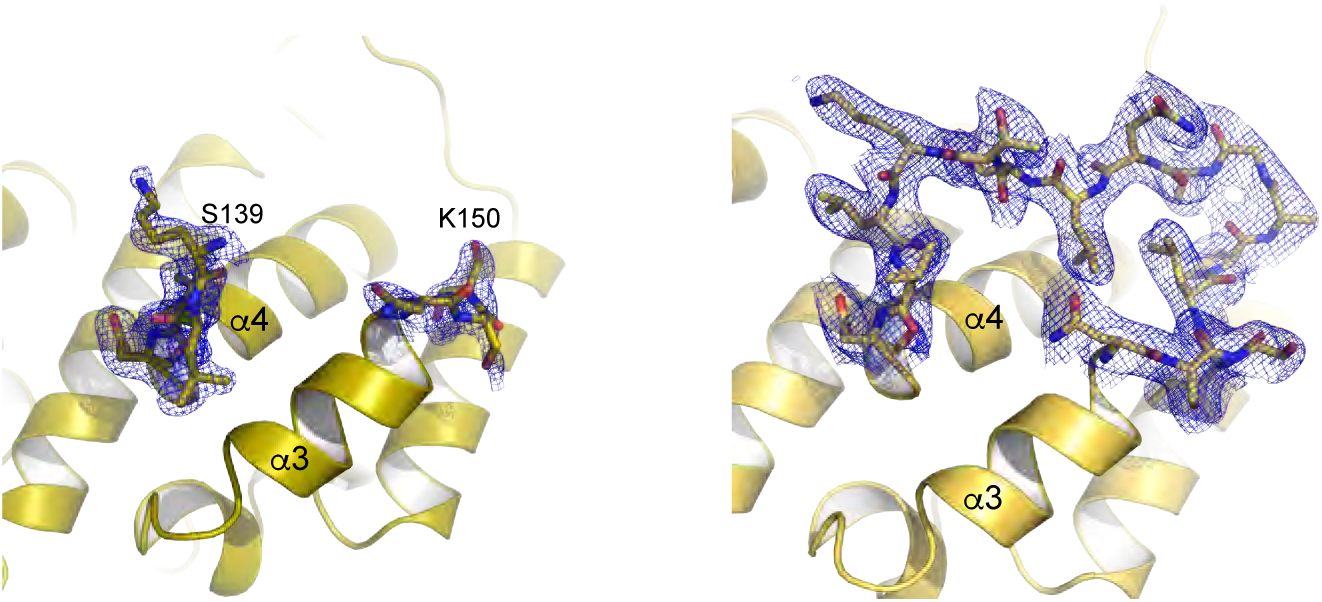
(Close view of the α3-α4 loop in apo (left panel) and holo (right panel) *Mt*SteB, with the corresponding electron densities contoured at 1 α.

**Figure S4.**
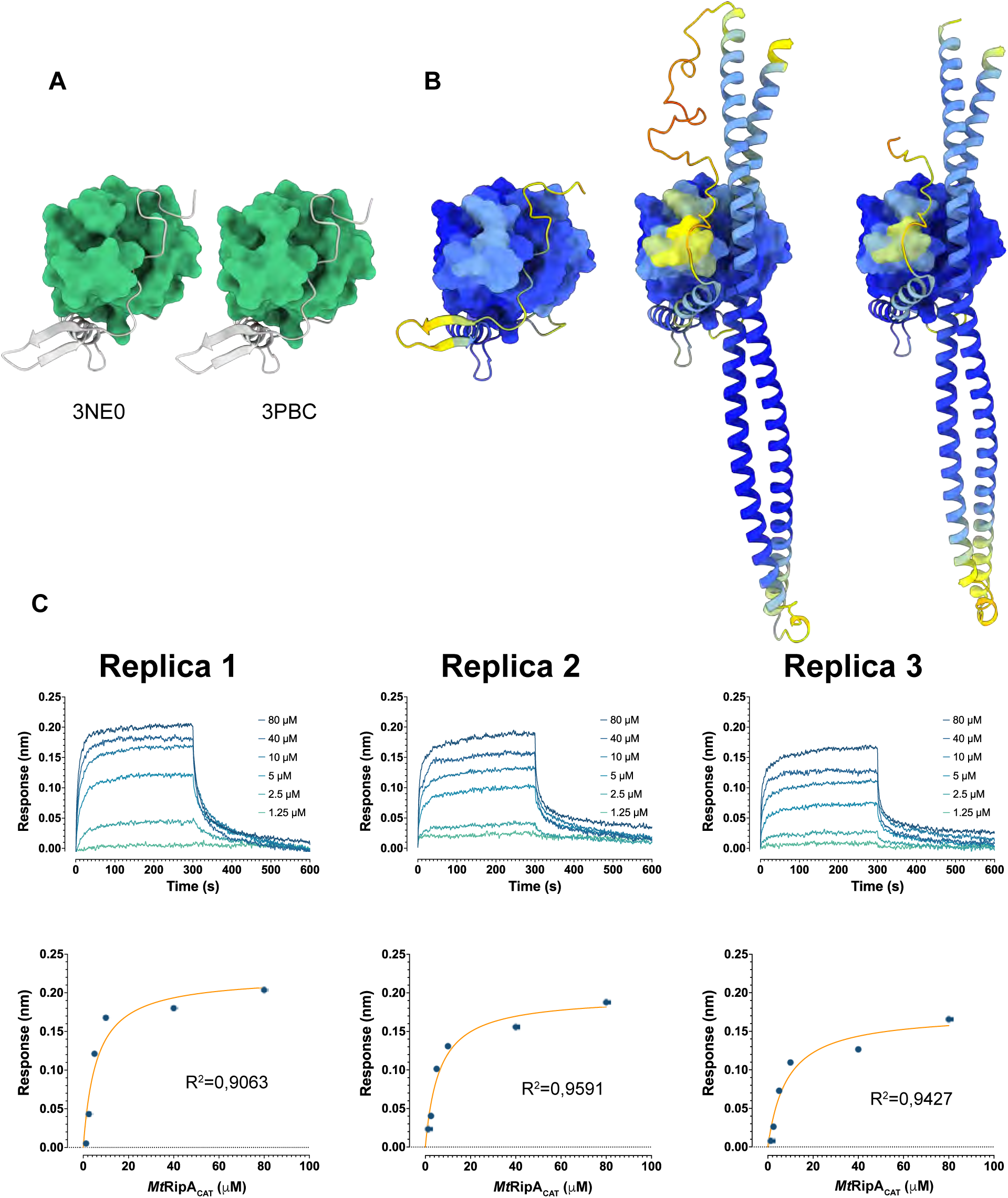
The *Mt*RipA catalytic domain is autoinhibited by the N-terminal CC. **(A)** Available crystal structures of truncated forms of the *Mt*RipA protein (residues 265-472), showing the NlpC/P60 catalytic domain occluded by the residue segment 264-279 (PDB codes 3NE0 and 3PBC). **(B)** The AF-predicted models of three different *Mt*RipA constructs color-coded by model confidence (left panel: construct comprising residues 265-472, pLDDT=91.5; central panel: construct comprising residues 40-472, pLDDT=86.6; and right panel: construct comprising residues 40-239+265-472, pLDDT=91.4) revealed the N-terminal CC, when present, bound to the active site, as seen in the structure of *Cglu* RipA (PDB code 8AUC, (1)). **(C)** Three independent BLI experiments showing the interaction profiles for the *Mt*RipA-*Mt*RipA_CAT_ interaction used to calculate the dissociation constant shown in Fig. 1F.

**Figure S5.**
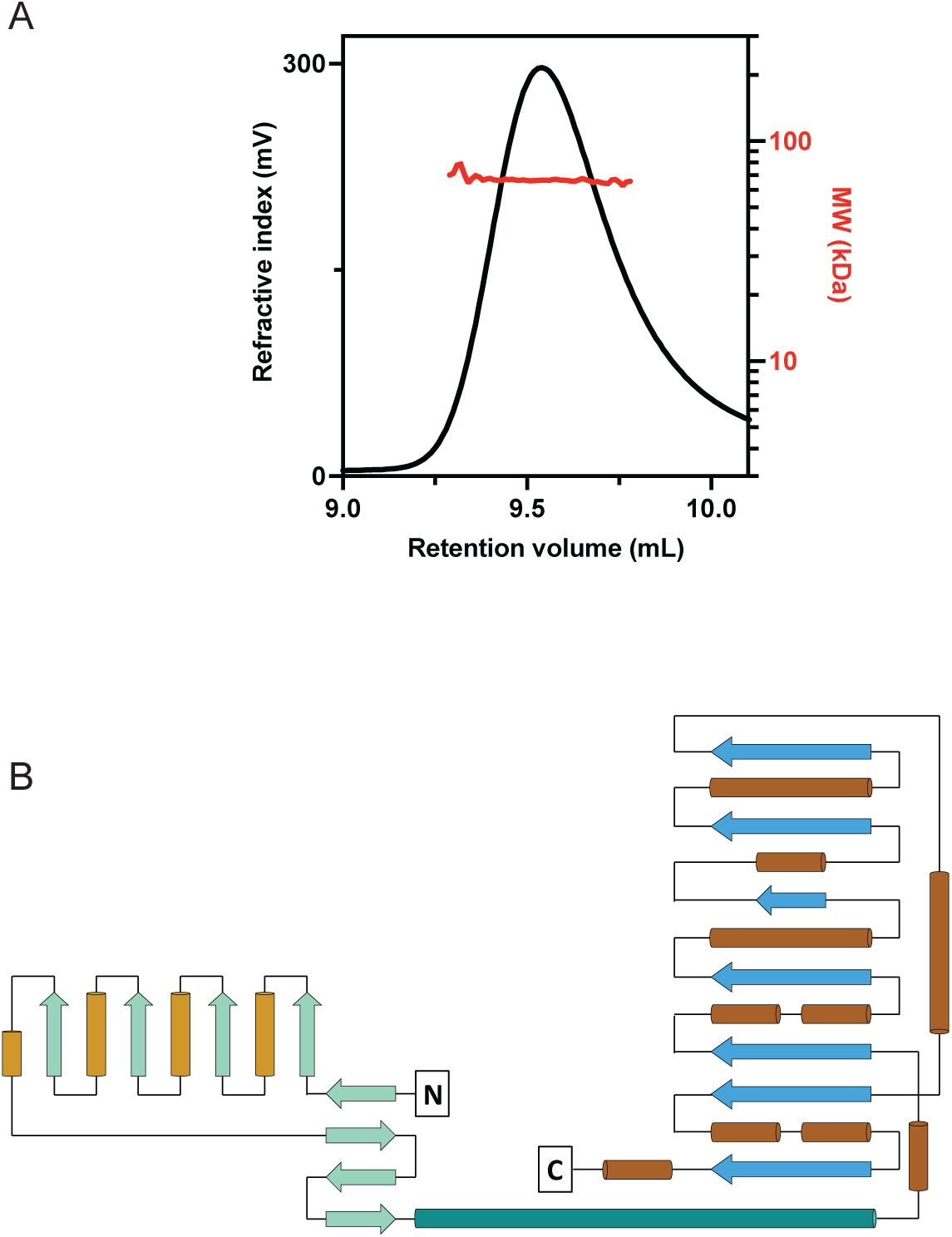
**(A)** SEC-SLS experiments show that *Mt*SteA elutes as a single peak with an apparent MW of 66 kDa, close to the predicted MW of the dimer (70 kDa). **(B)** *Mt*SteA secondary structure diagram.

**Figure S6.**
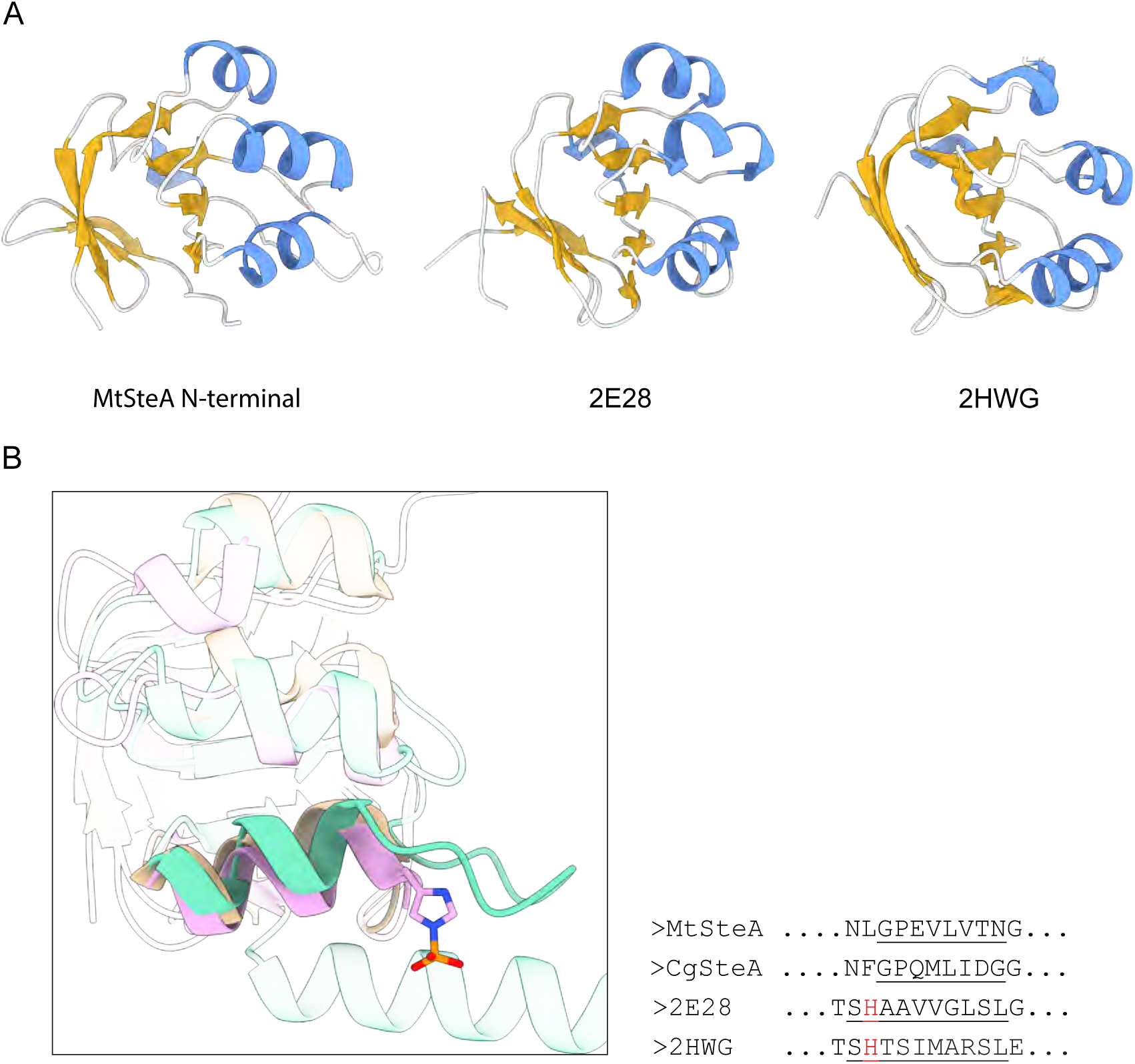
Structural similarity of the SteA N-terminal domain. (**A**) The N-terminal domain (left) can be superimposed with the phosphohistidine domains of pyruvate kinase (PDB code 2E28, (2)) with a rmsd of 1.051 Å for 48 equivalent Cα positions, and the phosphoenoylpyruvate:sugar phosphotransferase system (PDB code 2HWG, (3)) with a rmsd of 1.019 Å for 25 equivalent Cα positions. Structures are colored according to secondary structure (**B**) Close view of superposed SteA (green), 2E28 (tan) and 2HWG (pink). The (phospho)histidine residue is missing in *Mt*SteA. The partial sequence alignment of this region is shown at right.

**Figure S7.**
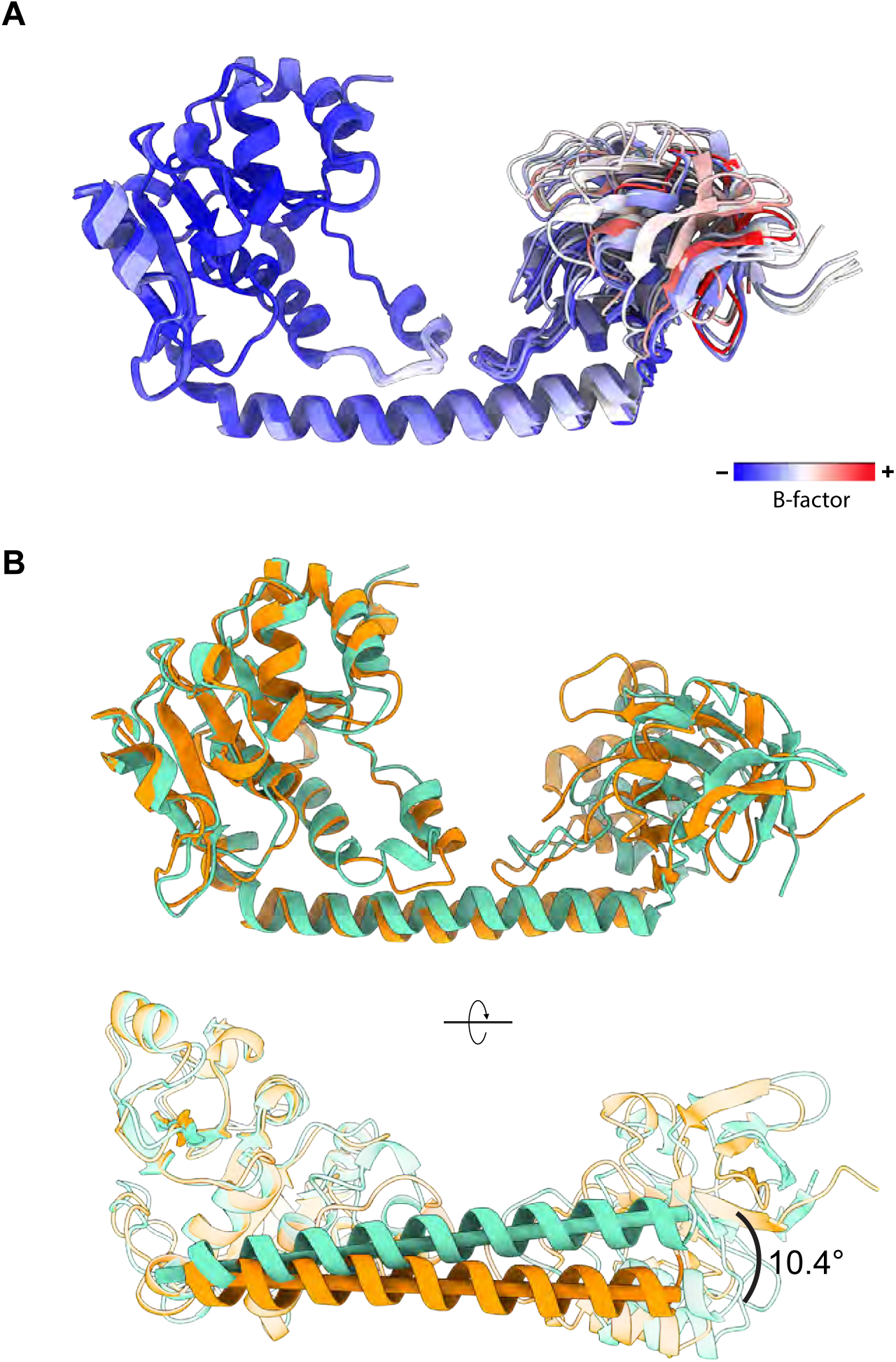
Structural flexibility of the SteA N-terminal domain. **(A)** Superposition of the 8 crystallographically independent molecules of *Cg*SteA color-coded according to B values. The highest B values are found for the N-terminal domain, on the right. **(B)** Two different views of the superposition between *Mt*SteA (green) and *Cg*SteA (orange, one molecule).

**Figure S8.**
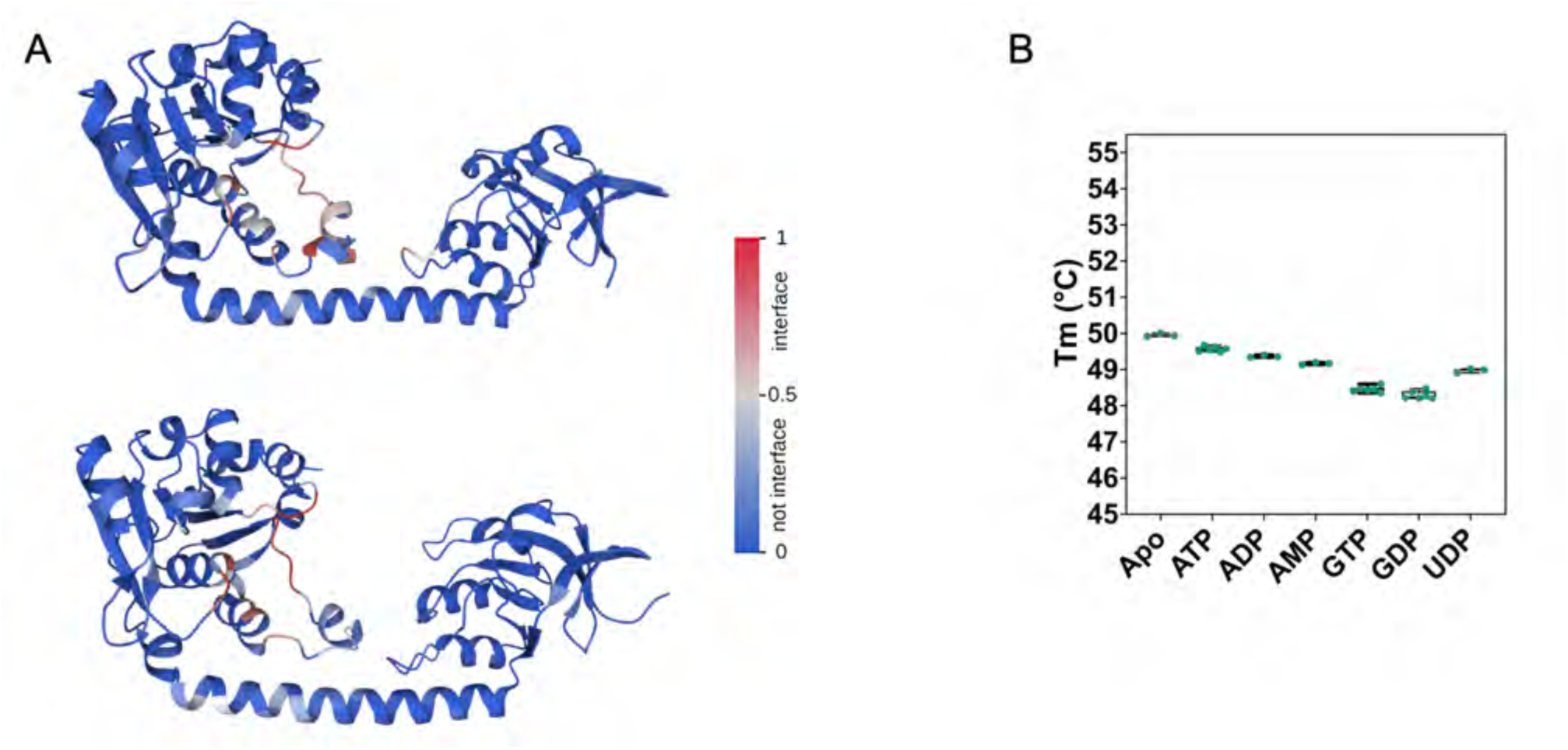
The phosphonucleotide-binding pocket of SteA. (**A**) The conserved binding pockets of *Mt*SteA (top) and *Cg*SteA (bottom) are both predicted to bind carbohydrate using PeSTo-Carbs, a Deep-Learning approach trained on protein-carbohydrate interfaces (4). (**B**) Binding of different phosphonucleotides to *Mt*SteA as assessed by nanoDSF were inconclusive. However, the same experiment carried out with *Cg*SteA (see Fig.3C, Main Text) indicated that the protein would bind UDP- or GDP-containing compounds.

**Figure S9.**
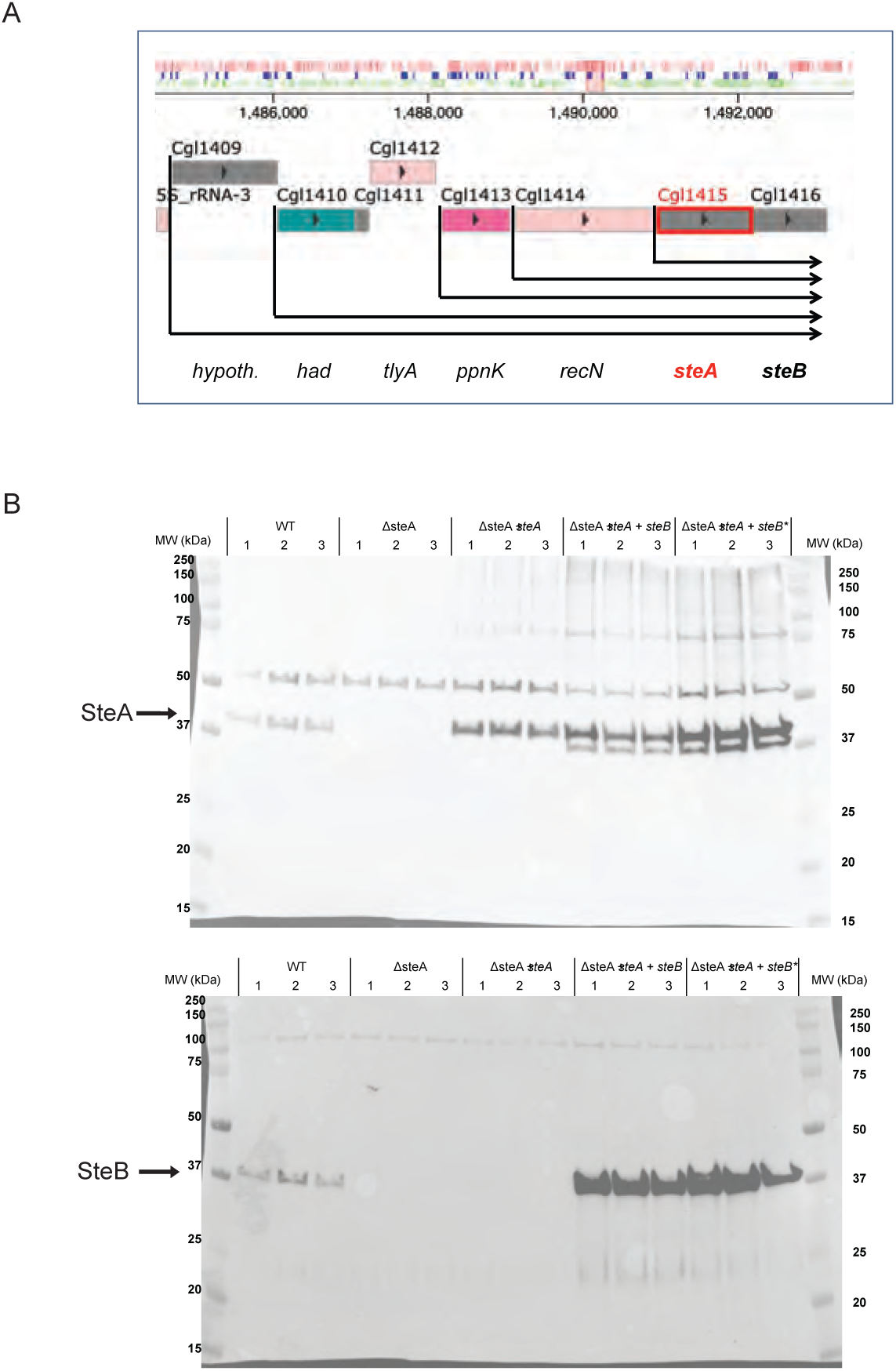
**(A)** Transcription start sites as described in (5) **(B)** Western Blots anti-SteA (top) and anti-SteB (bottom) of whole cell extracts (60 μg) from the different strains shown in Fig. 4. An arrow indicates the specific signal for *Cg*SteA and *Cg*SteB, respectively.

**Figure S10.**
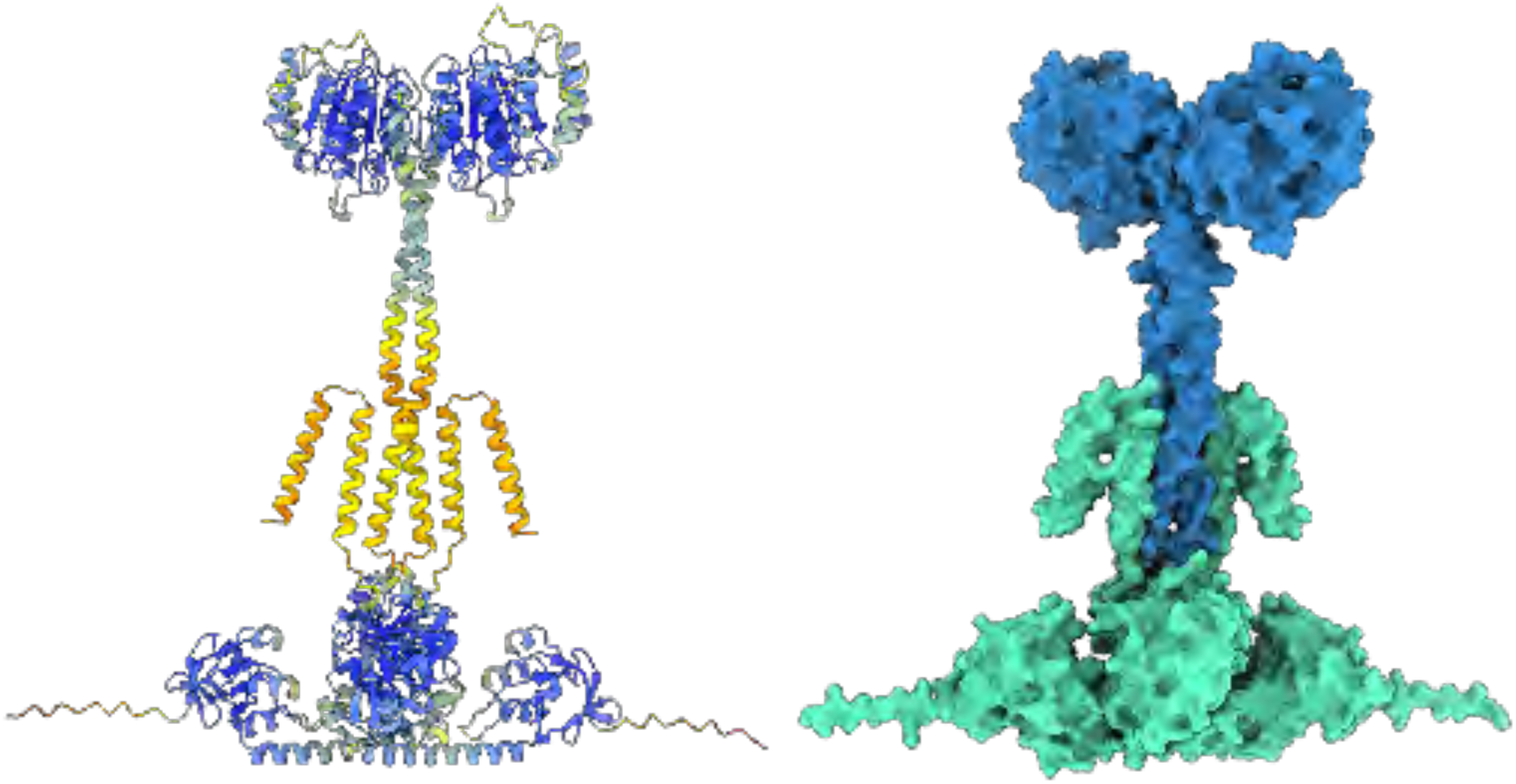
AlphaFold-predicted model of the heterotetramer (*Mt*SteA/*Mt*SteB)_2_ color-coded according to model confidence (left panel, plDDT=84) or protein organization (right panel), with the *Mt*SteA dimer shown in green and the *Mt*SteB dimer in blue.

**Figure S11.**
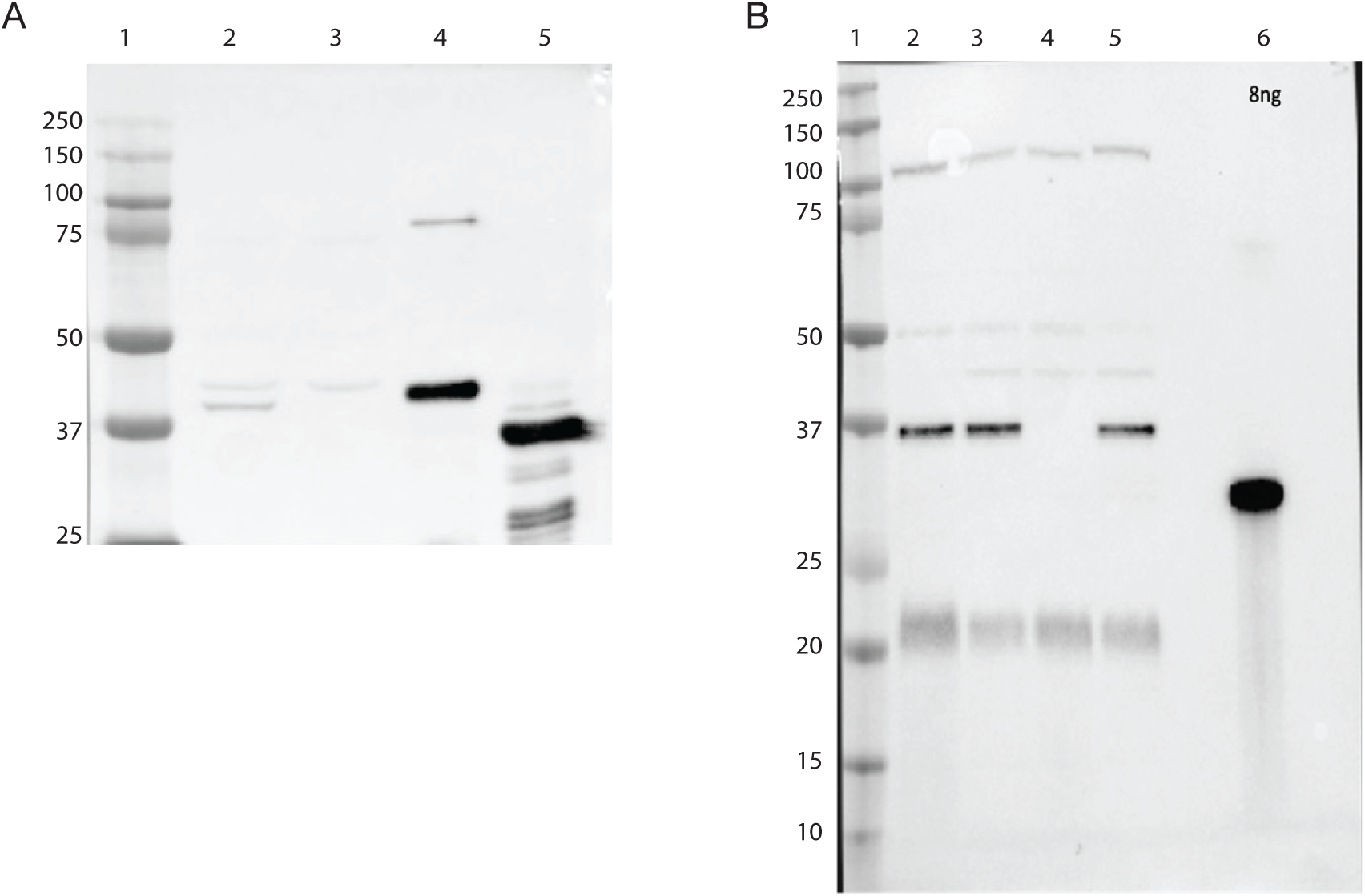
Antibody characterization. (**A**) Western Blot using purified anti-SteA antibody. Lane 1: molecular weight markers [kDa]; lane 2: total cell extract (120 μg) of *Cglu*; lane 3: total cell extract (120 μg) of *Cglu_λ1steAB*; lane 4: recombinant full-length His-SteA (0.1 μg); lane 5: recombinant soluble His-SteA (0.4 μg) (**B**) Western Blot using purified anti-SteB antibody. Lane 1: molecular weight markers, kDa; lane 2: total cell extract (120 μg) of *Cglu*; lane 3: total cell extract (120 μg) of *Cglu_empty plasmid*; lane 4: total cell extract (120 μg) of *Cglu_λ1steAB*; lane 5: total cell extract (120 μg) of *Cglu_λ1steAB + Scarlet-SteB* (note that in these conditions Scarlet-SteB is not expressed); lane 6: recombinant soluble His-SteB (8 ng).

**Supplementary Table S1.**
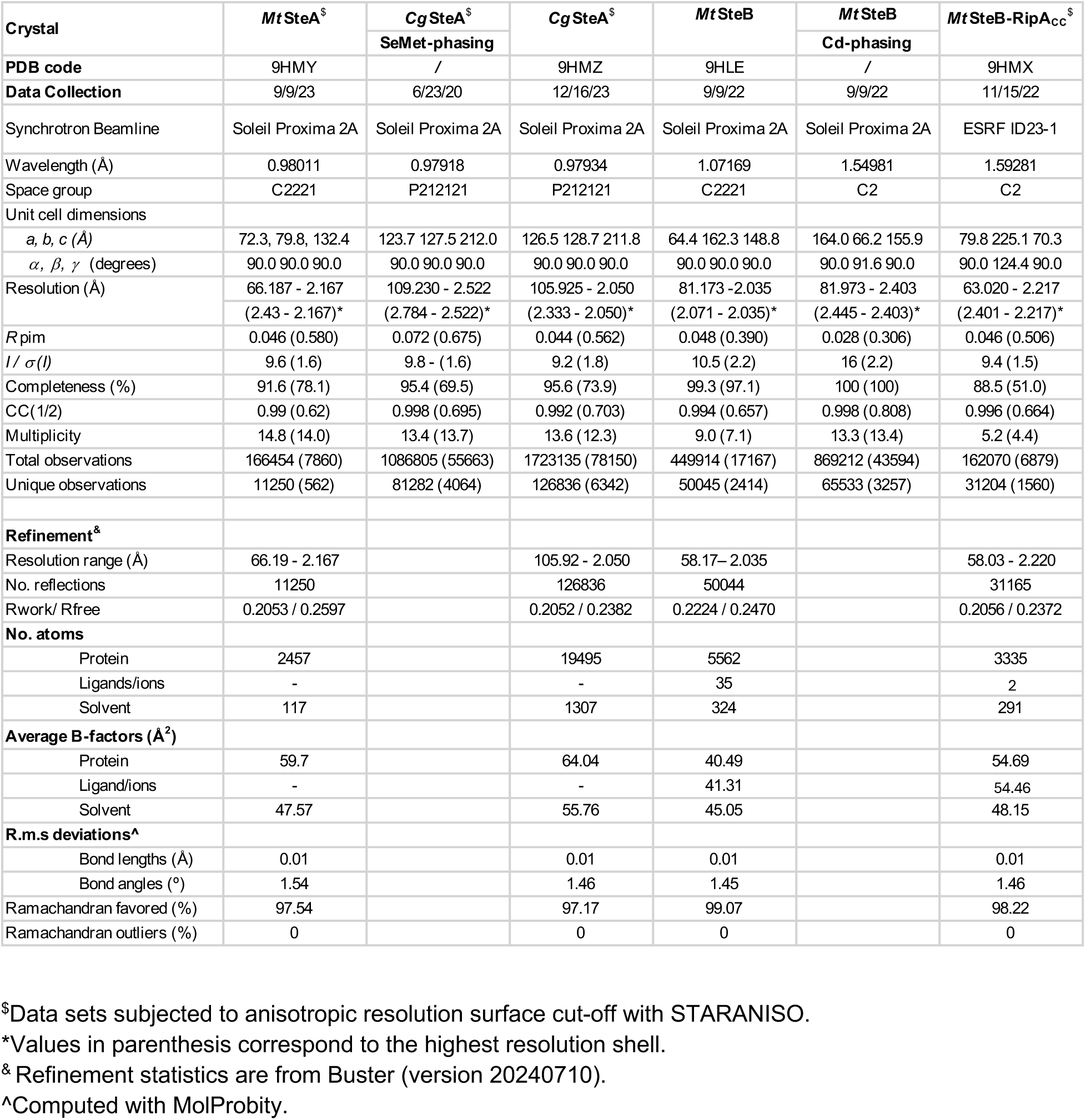
Crystallographic data collection and refinement statistics.

**Supplementary Table S2.**
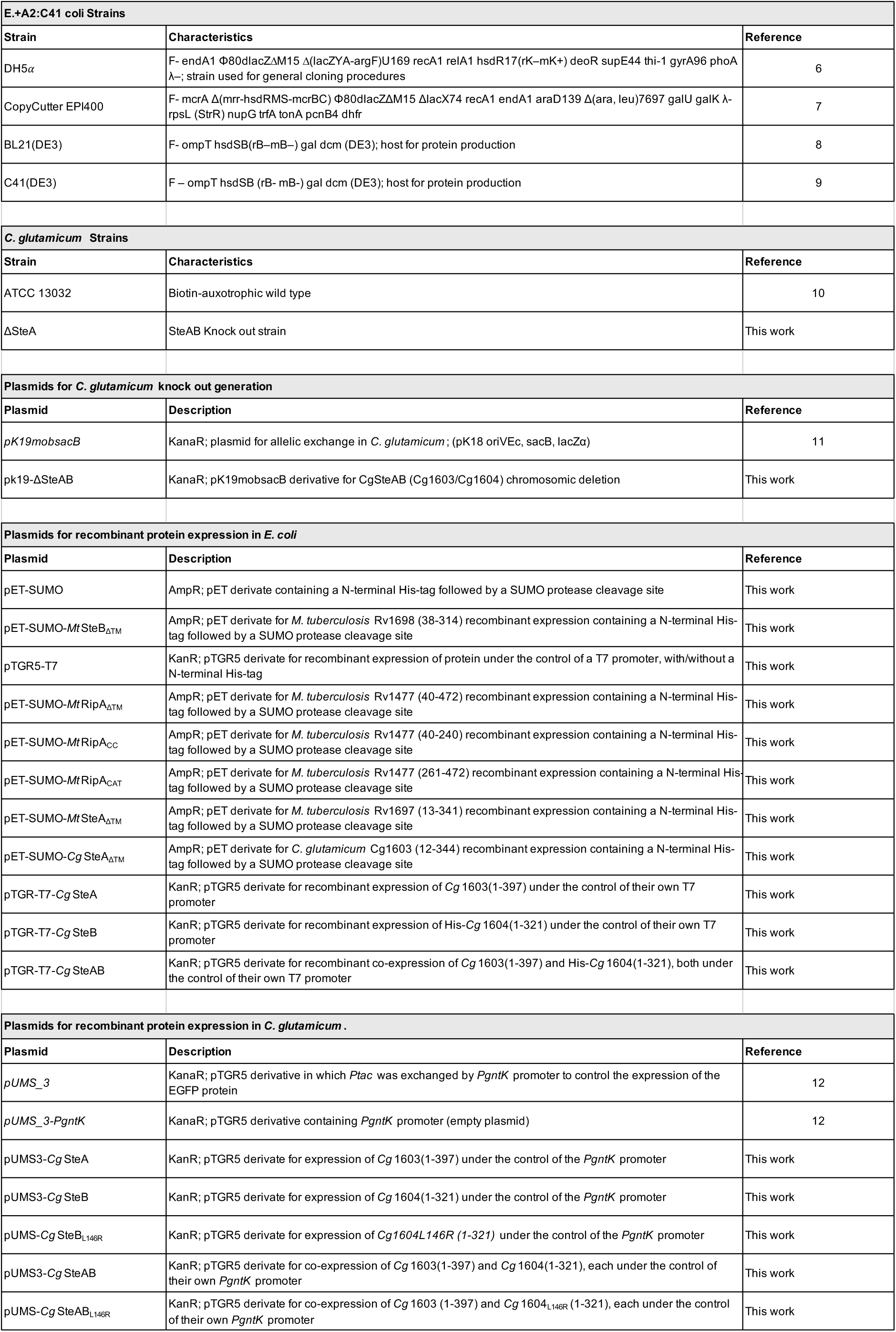
Plasmids and strains used in this study.

**Supplementary Table S3.**
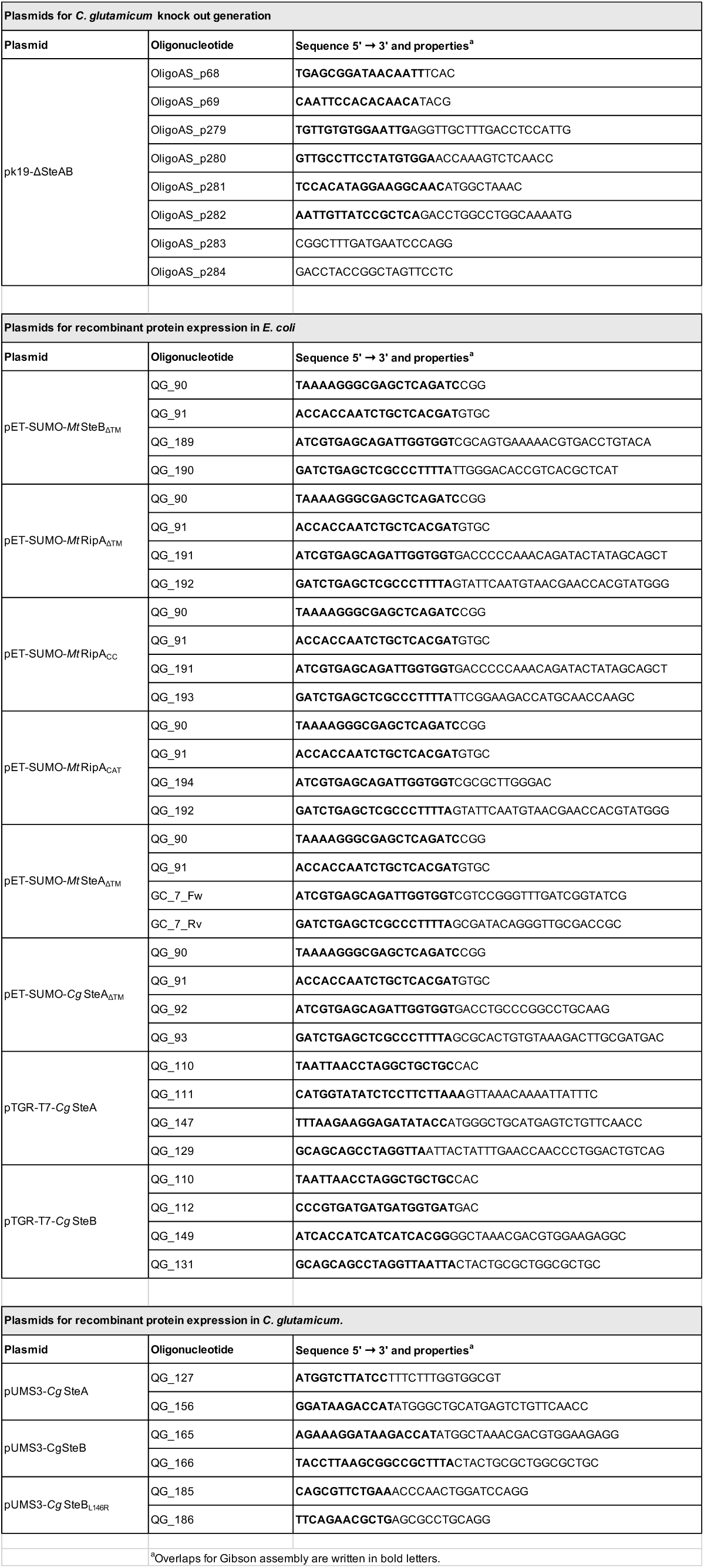
Oligonucleotide primers used in this study.

## References

1. A. J. F. Egan, R. M. Cleverley, K. Peters, R. J. Lewis, W. Vollmer, Regulation of bacterial cell wall growth. FEBS J. 32, 149 (2016).

2. X. Zhou, et al., Sequential assembly of the septal cell envelope prior to V snapping in *Corynebacterium glutamicum*. Nat Chem Biol 15, 221–231 (2019).

3. E. C. Hett, M. C. Chao, L. L. Deng, E. J. Rubin, A mycobacterial enzyme essential for cell division synergizes with resuscitation-promoting factor. PLoS Pathog 4, e1000001 (2008).

4. M. C. Chao, et al., Protein complexes and proteolytic activation of the cell wall hydrolase RipA regulate septal resolution in *Mycobacteria*. PLoS Pathog 9, e1003197 (2013).

5. C. Healy, A. Gouzy, S. Ehrt, Peptidoglycan hydrolases RipA and Ami1 are critical for replication and persistence of *Mycobacterium tuberculosis* in the host. Mbio 11, e03315–19 (2020).

6. L.-Y. Gao, M. Pak, R. Kish, K. Kajihara, E. J. Brown, A mycobacterial operon essential for virulence *in vivo* and invasion and intracellular persistence in macrophages. Infect Immun 74, 1757–1767 (2006).

7. Y. Tsuge, H. Ogino, H. Teramoto, M. Inui, H. Yukawa, Deletion of *cgR_*1596 and *cgR_*2070, encoding NlpC/P60 proteins, causes a defect in cell separation in *Corynebacterium glutamicum* R. J Bacteriol 190, 8204–8214 (2008).

8. L. Ott, et al., *Corynebacterium diphtheriae* invasion-associated protein (DIP1281) is involved in cell surface organization, adhesion and internalization in epithelial cells. Bmc Microbiol 10, 2 (2010).

9. E. C. Hett, et al., A partner for the resuscitation-promoting factors of *Mycobacterium tuberculosis*. Mol Microbiol 66, 658–668 (2007).

10. E. C. Hett, M. C. Chao, E. J. Rubin, Interaction and modulation of two antagonistic cell wall enzymes of mycobacteria. PLoS Pathog 6, e1001020 (2010).

11. H. Botella, et al., *Mycobacterium tuberculosis* protease MarP activates a peptidoglycan hydrolase during acid stress. EMBO J 36, 536–548 (2017).

12. Q. Gaday, et al., FtsEX-independent control of RipA-mediated cell separation in *Corynebacteriales*. Proc Natl Acad Sci USA 119, e2214599119 (2022).

13. H. C. Lim, et al., Identification of new components of the RipC-FtsEX cell separation pathway of *Corynebacterineae*. PLoS Genet 15, e1008284 (2019).

14. A. Siroy, et al., Rv1698 of *Mycobacterium tuberculosis* represents a new class of channel-forming outer membrane proteins. J Biol Chem 283, 17827–17837 (2008).

15. F. Wolschendorf, et al., Copper resistance is essential for virulence of *Mycobacterium tuberculosis*. Proc Natl Acad Sci USA 108, 1621–1626 (2011).

16. C. M. Sassetti, D. H. Boyd, E. J. Rubin, Genes required for mycobacterial growth defined by high density mutagenesis. Mol. Microbiol 48, 77–84 (2003).

17. J. E. Griffin, et al., High-resolution phenotypic profiling defines genes essential for mycobacterial growth and cholesterol catabolism. PLoS Pathog. 7, e1002251 (2011).

18. M. A. DeJesus, et al., Comprehensive essentiality analysis of the *Mycobacterium tuberculosis* genome via saturating transposon mutagenesis. Mbio 8, e02133–16 (2017).

19. Y. Minato, et al., Genome-wide assessment of *Mycobacterium tuberculosis* conditionally essential metabolic pathways. mSystems 4, 10.1128/msystems.00070-19 (2019).

20. A. Ruggiero, et al., Structure and functionalrRegulation of RipA, a mycobacterial enzyme essential for daughter cell separation. Structure 18, 1184–1190 (2010).

21. D. Böth, G. Schneider, R. Schnell, Peptidoglycan remodeling in *Mycobacterium tuberculosis*: comparison of structures and catalytic activities of RipA and RipB. J Mol Biol 413, 247–260 (2011).

22. E. M. Steiner, et al., The structure of the N-terminal module of the cell wall hydrolase RipA and its role in regulating catalytic activity. Proteins Struct Funct Bioinform 86, 912–923 (2018).

23. L. Holm, Dali server: structural unification of protein families. Nucleic Acids Res 50, W210–W215 (2022).

24. J.-Y. Liu, D. E. Timm, T. D. Hurley, Pyrithiamine as a substrate for thiamine pyrophosphokinase. J Biol Chem 281, 6601–6607 (2006).

25. C. P. C. Chiu, et al., Structural analysis of the sialyltransferase CstII from *Campylobacter jejuni* in complex with a substrate analog. Nat Struct Mol Biol 11, 163–170 (2004).

26. A. Schäfer, et al., Small mobilizable multi-purpose cloning vectors derived from the *Escherichia coli* plasmids pK18 and pK19: selection of defined deletions in the chromosome of *Corynebacterium glutamicum*. Genetics 145, 69–73 (1994).

27. K. Pfeifer-Sancar, A. Mentz, C. Rückert, J. Kalinowski, Comprehensive analysis of the *Corynebacterium glutamicum* transcriptome using an improved RNAseq technique. BMC Genomics 14, 888–888 (2013).

28. M. Jankute, J. A. G. Cox, J. Harrison, G. S. Besra, Assembly of the mycobacterial cell wall. Annu Rev Microbiol 69, 405–423 (2015).

29. A. Maitra, et al., Cell wall peptidoglycan in *Mycobacterium tuberculosis*: An Achilles’ heel for the TB-causing pathogen. FEMS Microbiol Rev 43, 548–575 (2019).

30. L. E. Ulrich, I. B. Zhulin, Four-helix bundle: a ubiquitous sensory module in prokaryotic signal transduction. Bioinformatics 21, iii45–iii48 (2005).

31. D. Albanesi, et al., Structural plasticity and catalysis regulation of a thermosensor histidine kinase. Proc Natl Acad Sci USA 106, 16185–16190 (2009).

32. F. Jacob-Dubuisson, A. Mechaly, J.-M. Betton, R. Antoine, Structural insights into the signalling mechanisms of two-component systems. Nat Rev Micro 16, 585–593 (2018).

33. A. Buschiazzo, F. Trajtenberg, Two-Component Sensing and Regulation: How Do Histidine Kinases Talk with Response Regulators at the Molecular Level? Annu Rev Microbiol 73, 1–22 (2019).

34. K. J. Wu, et al., Characterization of conserved and novel septal factors in *Mycobacterium smegmatis*. J Bacteriol 200 (2018).

35. J. Puffal, A. García-Heredia, K. C. Rahlwes, M. S. Siegrist, Y. S. Morita, Spatial control of cell envelope biosynthesis in mycobacteria. Pathog Dis 76, fty027 (2018).

36. J. L. Rowland, M. Niederweis, Resistance mechanisms of *Mycobacterium tuberculosis* against phagosomal copper overload. Tuberculosis 92, 202–210 (2012).

37. R. A. Slayden, et al., Updating and curating metabolic pathways of TB. Tuberculosis 93, 47–59 (2013).

38. R. Wang, et al., Persistent *Mycobacterium tuberculosis* infection in mice requires PerM for successful cell division. Elife 8, e49570 (2019).

39. N. Goodsmith, et al., Disruption of a *M. tuberculosis* membrane protein causes a magnesium-dependent cell division defect and failure to persist in mice. PLoS Pathog. 11, e1004645 (2015).

40. Y. Ju, et al., The gene MAB_2362 is responsible for intrinsic resistance to various drugs and virulence in *Mycobacterium abscessus* by regulating cell division. Antimicrob Agents Chemother 69, e00433–24 (2024).

41. J. S. Philalay, C. O. Palermo, K. A. Hauge, T. R. Rustad, G. A. Cangelosi, Genes required for intrinsic multidrug resistance in *Mycobacterium avium*. Antimicrob Agents Chemother 48, 3412–3418 (2004).

42. M. A. Mir, H. S. Rajeswari, U. Veeraraghavan, P. Ajitkumar, Molecular characterisation of ABC transporter type FtsE and FtsX proteins of *Mycobacterium tuberculosis*. Arch Microbiol 185, 147–158 (2006).

43. D. Mavrici, et al., *Mycobacterium tuberculosis* FtsX extracellular domain activates the peptidoglycan hydrolase, RipC. Proc Natl Acad Sci USA 111, 8037–8042 (2014).

44. K. J. Kieser, E. J. Rubin, How sisters grow apart: mycobacterial growth and division. Nat Rev Microbiol 12, 550–562 (2014).

45. J. Li, et al., Regulation of the cell division hydrolase RipC by the FtsEX system in *Mycobacterium tuberculosis*. Nat Commun 14, 7999 (2023).

46. N. R. Thanky, D. B. Young, B. D. Robertson, Unusual features of the cell cycle in mycobacteria: Polar-restricted growth and the snapping-model of cell division. Tuberculosis 87, 231–236 (2007).

47. X. Zhou, D. K. Halladin, J. A. Theriot, Fast mechanically driven daughter cell separation is widespread in *Actinobacteria*. mBio 7, e00952–16 (2016).

48. P. D. Odermatt, et al., Overlapping and essential roles for molecular and mechanical mechanisms in mycobacterial cell division. Nat Phys 16, 57–62 (2020).

49. B. Miroux, J. E. Walker, Over-production of proteins in *Escherichia coli*: mutant hosts that allow synthesis of some membrane proteins and globular proteins at high levels. J Mol Biol 260, 289–298 (1996).

50. C. Keilhauer, L. Eggeling, H. Sahm, Isoleucine synthesis in *Corynebacterium glutamicum*: molecular analysis of the *ilvB-ilvN-ilvC* operon. J Bacteriol 175, 5595–5603 (1993).

51. P. Ravasi, S. Peiru, H. Gramajo, H. G. Menzella, Design and testing of a synthetic biology framework for genetic engineering of *Corynebacterium glutamicum*. Microb Cell Fact 11, 147 (2012).

52. F. W. Studier, Protein production by auto-induction in high-density shaking cultures. Protein Expr Purif 41, 207–234 (2005).

53. P. Weber, et al., High-throughput crystallization pipeline at the crystallography core facility of the Institut Pasteur. Molecules 24, 4451 (2019).

54. W. Kabsch, XDS. Acta Crystallogr Sect D Biol Crystallogr 66, 125–32 (2010).

55. M. D. Winn, et al., Overview of the CCP4 suite and current developments. Acta Crystallogr Sect D Biol Crystallogr 67, 235–42 (2011).

56. C. Vonrhein, et al., Data processing and analysis with the autoPROC toolbox. Acta Crystallogr Sect D Biol Crystallogr 67, 293–302 (2011).

57. I. J. Tickle, et al., STARANISO. Cambridge, United Kingdom: Global Phasing Ltd. (2017).

58. P. Skubák, et al., A new MR-SAD algorithm for the automatic building of protein models from low-resolution X-ray data and a poor starting model. IUCr J 5, 166–171 (2018).

59. P. Emsley, K. Cowtan, Coot: model-building tools for molecular graphics. Acta Crystallogr Sect D Biol Crystallogr 60, 2126–2132 (2004).

60. D. Liebschner, et al., Macromolecular structure determination using X-rays, neutrons and electrons: recent developments in Phenix. Acta Crystallogr Sect D Biol Crystallogr 75, 861–877 (2019).

61. O. S. Smart, et al., Exploiting structure similarity in refinement: automated NCS and target-structure restraints in BUSTER. Acta Crystallogr Sect D Biol Crystallogr 68, 368–380 (2012).

62. T. D. Goddard, et al., UCSF ChimeraX: Meeting modern challenges in visualization and analysis. Protein Sci 27, 14–25 (2018).

63. J. Schindelin, et al., Fiji: an open-source platform for biological-image analysis. Nat Methods 9, 676–682 (2012).

64. K. J. Cutler, et al., Omnipose: a high-precision morphology-independent solution for bacterial cell segmentation. Nat Methods 19, 1438–1448 (2022).

65. A. Ducret, E. M. Quardokus, Y. V. Brun, MicrobeJ, a tool for high throughput bacterial cell detection and quantitative analysis. Nat Microbiol 1, 16077 (2016).

66. M. Martinez, et al., Eukaryotic-like gephyrin and cognate membrane receptor coordinate corynebacterial cell division and polar elongation. Nat Microbiol 8, 1896–1910 (2023).

67. A. Sogues, et al., Essential dynamic interdependence of FtsZ and SepF for Z-ring and septum formation in *Corynebacterium glutamicum*. Nat Commun 11, 1641 (2020).

68. G. Madeo, C. Savojardo, M. Manfredi, P. L. Martelli, R. Casadio, CoCoNat: a novel method based on deep learning for coiled-coil prediction. Bioinformatics 39, btad495 (2023).

69. H. Ashkenazy, et al., ConSurf 2016: an improved methodology to estimate and visualize evolutionary conservation in macromolecules. Nucleic Acids Res. 44, W344–W350 (2016).

70. B. Zuber, et al., Direct visualization of the outer membrane of *Mycobacteria* and *Corynebacteria* in their native state. J Bacteriol 190, 5672–5680 (2008).

71. B. Isbilir, A. Yeates, V. Alva, T. A. M. Bharat, Mapping the ultrastructural topology of the corynebacterial cell surface. bioRxiv 2024.09.05.611374 (2024).

## SI References

1. Q. Gaday, et al., FtsEX-independent control of RipA-mediated cell separation in *Corynebacteriales*. Proc Natl Acad Sci USA 119, e2214599119 (2022).

2. A. Teplyakov, et al., Structure of phosphorylated enzyme I, the phosphoenolpyruvate:sugar phosphotransferase system sugar translocation signal protein. Proc Natl Acad Sci 103, 16218–16223 (2006).

3. K. Suzuki, S. Ito, A. Shimizu-Ibuka, H. Sakai, Crystal structure of pyruvate kinase from *Geobacillus stearothermophilus*. J Biochem 144, 305–312 (2008).

4. K. Pfeifer-Sancar, A. Mentz, C. Rückert, J. Kalinowski, Comprehensive analysis of the *Corynebacterium glutamicum* transcriptome using an improved RNAseq technique. BMC Genomics 14, 888–888 (2013).

5. P. Bibekar, L. Krapp, M. Dal Peraro, PeSTo-Carbs: geometric deep learning for prediction of protein-carbohydrate binding interfaces. J Chem Theory Comput 20, 2985–91 (2024).

6. D. Hanahan, Studies on transformation of *Escherichia coli* with plasmids. J Mol Biol 166, 557–80 (1983)

7. D. Hasking. Epicentre Forum 11, 6 (2004).

8. F.W. Studier, B.A. Moffatt, User of bacteriophage T7 RNA polymerase to direct selective high-level expression of cloned genes. J Mol Biol 189, 113–30 (1986).

9. B. Miroux, J. E. Walker, Over-production of proteins in *Escherichia coli*: mutant hosts that allow synthesis of some membrane proteins and globular proteins at high levels. J Mol Biol 260, 289–298 (1996).

10. S. Kinoshita, S. Udaka, M. Shimono, Studies on the amino acid fermentation. J Gen Appl Microbiol 3, 193–205 (1957)

11. A. Schäfer, et al., Small mobilizable multi-purpose cloning vectors derived from the *Escherichia coli* plasmids pK18 and pK19: selection of defined deletions in the chromosome of *Corynebacterium glutamicum*. Genetics 145, 69–73 (1994).

12. M. Martinez, et al., Eukaryotic-like gephyrin and cognate membrane receptor coordinate corynebacterial cell division and polar elongation. Nat Microbiol 8, 1896–1910 (2023).

